# Naturally acquired IgG responses to *Plasmodium falciparum* do not target the conserved termini of the malaria vaccine candidate merozoite surface protein 2

**DOI:** 10.1101/2024.09.24.614082

**Authors:** Julia Zerebinski, Lucille Margerie, Nan Sophia Han, Maximilian Moll, Matias Ritvos, Peter Jahnmatz, Niklas Ahlborg, Billy Ngasala, Ingegärd Rooth, Ronald Sjöberg, Christopher Sundling, Victor Yman, Anna Färnert, David Plaza

**Affiliations:** Karolinska Institutet, Division of Infectious Diseases, Department of Medicine Solna and Center for Molecular Medicine, Stockholm, Sweden; Karolinska University Hospital, Department of Infectious Diseases, Stockholm, Sweden; University of Bonn, University Hospital of Bonn, Medical Clinic III for Oncology, Hematology, Immuno-Oncology and Rheumatology, Bonn, Germany; MabTech, Nacka, Sweden; Muhimbili University of Health and Allied Sciences, Department of Parasitology and Medical Entomology, Dar es Salaam, Tanzania; SciLifeLab, Autoimmunity and Serology Profiling Unit, Solna, Sweden; Södersjukhuset, Department of Infectious Diseases, Stockholm, Sweden; Institut Pasteur Paris, Department of Global Health, Infectious Disease Epidemiology & Analytics Unit, Paris, France

**Keywords:** Malaria vaccine, merozoite, polymorphic antigens, immune evasion, structural heterogeneity

## Abstract

Malaria remains a significant burden, and a fully protective vaccine against *Plasmodium falciparum* is critical for reducing morbidity and mortality. Antibody responses against the blood-stage antigen merozoite surface protein 2 (MSP2) are associated with protection from *P. falciparum* malaria, but its extensive polymorphism is a barrier to its development as a vaccine candidate. New tools, such as long-read sequencing and accurate protein structure modelling allow us to more easily study the genetic diversity and immune responses towards antigens from clinical isolates. This study sought to better understand naturally acquired MSP2-specific antibody responses. IgG responses against recombinantly expressed full-length, central polymorphic regions, and peptides derived from the conserved termini of MSP2 variants sequenced from patient isolates, were tested in plasma from travelers with recent, acute malaria and from individuals living in an endemic area of Tanzania. IgG responses towards full MSP2 and truncated MSP2 antigens were variant specific. IgG antibodies in the plasma of first-time infected or previously exposed travelers did not recognize the conserved termini of expressed MSP2 variants by ELISA, but they bound 13- amino acid long linear epitopes from the termini in a custom-made peptide array. Alphafold3 modelling suggests extensive structural heterogeneity in the conserved termini upon antigen oligomerization. IgG from individuals living in an endemic region, many who were asymptomatically infected, did not recognize the conserved termini by ELISA. Our results suggest that responses to the variable regions are important for the development of naturally acquired immunity towards MSP2.

## 1 Introduction

Polymorphic antigens are challenging vaccine targets and need to be better understood in the context of human immunity (1). The *Plasmodium falciparum* merozoite surface protein 2 (MSP2) is no exception (2). Of the five human *Plasmodium* parasite species, *P. falciparum* accounts for over 95% of cases and deaths globally (3). The World Health Organization estimated 249 million malaria cases in 2022 which resulted in 608,000 deaths (3). An efficacious vaccine against *P. falciparum* would be a key tool to decrease morbidity and mortality. The WHO-recommended RTS,S/AS01 and R21/MatrixM vaccines target the pre-erythrocytic stage circumsporozoite surface protein (CSP) (4–7). This single-stage and single-antigen approach represents a challenge for successful vaccination as parasite escape leads to hepatocyte and blood infection (8,9). A blood-stage vaccine, or a multivalent vaccine including both blood-stage and pre-erythrocytic stage antigens, could prevent clinical malaria and death (10,11).

MSP2 is an abundant, intrinsically disordered protein covalently bound to the merozoite membrane by a glycosylphosphatidylinositol (GPI) anchor (12). The *msp2* gene is made up of two conserved termini flanking a highly polymorphic central domain (13,14), consisting of repetitive motifs flanked by dimorphic repeat sequences which divide MSP2 variants into two allelic families: FC27 and 3D7 (IC) (2,13). There is conflicting evidence on the importance of MSP2 to the invasion of erythrocytes and the relative fitness of the parasite (15), and its function remains unknown. The high degree of polymorphism of MSP2 likely allows the parasite to escape immune recognition in the blood stream (16) and is believed to slow development of memory B and T cell responses. Its extensive diversity is also a significant barrier to its further development as a vaccine candidate.

The diversity of MSP2 also makes it a prime example for the study of human antibody responses to polymorphic proteins, especially those under immune selection (17,18). Natural *P. falciparum* infection induces antibodies against merozoite surface proteins, which are associated with reduced risk of clinical malaria and morbidity (19–22). Mouse monoclonal antibodies against MSP2 have previously been shown to target the central variable region, central repeats, and conserved termini (23). Studies using mice immunized with MSP2 showed immunogenicity of the conserved termini (24), yet in vaccinated children antibodies against single antigenic variants were not sufficient for protection (25). Instead, responses against multiple antigen variants, resulting in a cumulative protective effect, are likely necessary for clinical immunity (19,26). Such multivalent vaccines are already in development or approved for influenza (27) and human papillomavirus (28). With the aim to better understand naturally acquired antibody responses towards antigenic variants of MSP2, we investigated the binding of IgG in plasma from individuals with different levels of exposure, to recombinantly expressed full length MSP2, the central polymorphic region of MSP2, and conserved terminus peptides, sequenced from parasite isolates.

### 2 Methods

#### 2.1 Study Participants

Adults diagnosed with *P. falciparum* malaria after traveling to malaria-endemic areas were treated at Karolinska University Hospital (Stockholm, Sweden) and included in a prospective cohort as described previously (29,30). The study included travelers who experienced their primary *P. falciparum* malaria episode (n=13) (called “primary infected”), as well as those with a reported earlier *P. falciparum* infection or lived in a malaria-endemic area for at least one year, who were called “previously exposed” (n=23). Adults and children living in Nyamisati, a rural fishing village on the coast of Tanzania, participating in cross-sectional surveys in 1994 and in 1999 at the start of the rainy season were also studied (21,31,32). Samples were excluded if the participant was treated with antimalarials within 4 weeks prior to or 1 week after survey. At the time of sampling, malaria transmission was perennial with seasonal variation due to rainy seasons, with high parasite prevalence of 73.9% in 1994 and 66.3% in 1999 by PCR (32). For both cohorts, venous blood samples were collected in EDTA, frozen and stored separately as plasma and packed red blood cells at –80LJC.

#### 2.2 Ethics

Studies of *P. falciparum* antigen diversity and antibody responses in travelers and in the Tanzania cohort were approved by the Stockholm Region Ethical Committee and the Swedish Ethical Review Authority. The study was also approved by the Ethical Review board of the National Institute for Medical Research in Tanzania. Informed consent was obtained from all study participants or their guardians.

#### 2.3 PCR amplification and long-read amplicon sequencing with *in silico* genotyping

Nested amplification and high-fidelity long-read sequencing of the complete open reading frame of *msp2* was performed as described previously (33) on *P. falciparum*-positive samples using the Applied Biosystems SimpliAmp™ Thermal Cycler (Thermo Fisher Scientific). All primers are detailed in **Table S1**. Briefly, a primary PCR reaction amplified the complete open reading frame of the *msp2* gene under the following conditions: 98°C for 30 seconds, 40 cycles of: 98°C for 10 seconds, 58.1°C for 30 seconds, and 72°C for 40 seconds, followed by 72°C for 5 minutes. The primary PCR reaction included 1x Phusion HF Buffer, 6U Phusion Hot Start DNA Polymerase (Fischer Scientific), 200µM dNTP mix, and 0.5µM each forward and reverse primers (Fischer Scientific) and 1μL purified DNA per 15μL total volume reaction. A second nested PCR reaction targeted both allelic families of *msp2* by amplifying the region directly downstream and upstream of the annealing sites for msp2_fw and msp2_rv, respectively, under the following conditions: 98°C for 30 seconds, followed by 6 cycles of 98°C for 10 seconds, 50°C for 30 seconds, and 72°C for 40 seconds; 35 cycles of 98°C for 10 seconds and 72°C for 40 seconds, and followed by 72°C for 5 minutes. The secondary reaction included 1x Phusion HF Buffer, 20U Phusion Hot Start II DNA Polymerase (Fischer Scientific), 0.5µM of each forward and reverse primers (Fischer Scientific), plus 1µl product from the primary reaction in 50μL of final volume. Primers for the secondary reaction included unique barcodes which, in combination, allowed the identification of specific isolates for every sequencing read. PCR products from up to 12 reactions were pooled and cleaned using QIAquick PCR Purification Kit (QIAGEN) according to manufacturer instructions. A 1% agarose gel was used to identify successful amplification in samples and positive controls after both PCR reactions and product clean-up, as well as the lack of amplification in the individual water (negative) controls included in the library. Circular consensus sequencing (CCS) was performed using SMRTcell™ 1M on a PacBio Sequell I instrument with a Sequel 3.0 polymerase. An analysis pipeline for size fragment genotyping using CCS reads was developed previously (33). Briefly, an *in silico* PCR protocol was created whereby IC and FC27 primer sequences used for size fragment analysis (34), were run as Blastn queries against the sequencing dataset (**Table S1**) and filtered using a false positivity rate of 0.01 for both size and sequence calling.

#### 2.4 Expression of recombinant MSP2 antigens

Two FC27 and two IC *msp2* sequences from *P. falciparum* isolates in travelers were selected for recombinant expression. Variants were selected if at least 25 reads were found to support the sequence in the library, to exclude potential sequencing artefacts from the analysis. The four selected MSP2 sequences were aligned (**Figure S2A**) with reference strains 3D7 and Dd2. Consensus score for the four sequences was calculated in Jalview using normalized Valdar consensus and plotted for each residue of the six aligned sequences **(Figure S2B)**. From each of the four expressed proteins, two recombinant constructs were produced: one full-length construct, called “Full”, and one construct lacking the conserved N- and C-terminus regions, called “Truncated” or “Trunc.”. All proteins had native signal peptides and GPI-anchor signals removed. N-glycosylation sites were predicted using the NetNGlyc 1.0 Server (35), and predicted glycosylation motifs were modified by replacing Serines or Threonines with Alanines. A signal peptide from mouse IgG kappa chain was added to the N-terminus of all sequences. All constructs were engineered to include a C-tag (EPEA) at the N-terminus just downstream from the added signal peptide. In addition, Twin-Strep (SAWSHPQFEKGGGSGGGSGGSAWSHPQFEK), GAL (YPGQAPPGAYPGQAPPGA), WASP (CPDYRPYDWASPDYRD) and TRAP (DDFLSQQPERPRDVKLA) tags were added to the C-terminus of MSP2-FC27 1, MSP2-IC 1, MSP2-IC 2, and MSP2-FC27 2, respectively, to facilitate downstream immunofluorescence experiments. Tagged, codon-optimized sequences were added to pcDNA3.1/Zeo(-) plasmids by GeneScript (Piscataway, NJ, USA). Transfection and recombinant protein expression in HEK293T/17 (RRID:CVCL_1926) cells were performed as previously described (36). Finally, two synthetic peptides were produced for the conserved N- (YSNTFINNAYNMAIRRSM) and C- (AAPENKGTGQHGHMHGSRNNHPQNTADSQKECTDGNKENCGAATSLL) termini of MSP2 using the microwave-assisted PepPower™ peptide synthesis platform (GenScript, Piscataway, NJ, USA). Oligomerization of all recombinant proteins and synthetic peptides was assessed by SDS-PAGE after boiling (95LJC) protein or peptide aliquots in Laemmli buffer for 5 min in the presence or absence of dithiothreitol (DTT), respectively.

#### 2.5 IgG ELISA

Anti-MSP2 IgG ELISA was performed as previously described with minor modifications (37). Briefly, individual wells of 96-well flat bottom microplates (Corning Inc.) were coated with carbonate coating buffer (pH 9.3; 35mM NaHCO_3_, 15mM Na_2_CO_3_) containing 28.02nM of antigen. Plates were incubated overnight at 4°C before washing and blocking for 5h at room temperature with 1% Bovine Serum Albumin in PBS-Tween (PBS-0.05% Tween 20). Travelers’ samples were diluted 1:2000 while samples from Tanzania donors were diluted 1:4000 in assay buffer due to the higher concentration of antibodies in the latter. Negative controls of unexposed plasma pools were diluted 1:2000 for testing of Travelers samples and 1:4000 for Tanzania samples. Consistency in the absorbance of a hyperimmune control pool of Kenyan adult plasmas, diluted 1:4000, was used on every plate to monitor the reproducibility of the absorbance measured for different plates. Wells were incubated overnight at 4°C with test plasma. Plates were then washed and incubated for 3 hours at room temperature with HRP-conjugated rabbit anti-human IgG (Agilent) at 1:5000 dilution followed by extensive washing and incubation with TMB substrate (TMB Stabilized Chromogen, Invitrogen) for 15 minutes. The reaction was stopped with 0.18M H_2_SO_4_. Optical density (OD) was measured at 450nm using a microplate spectrophotometer (Multiskan SkyHigh, Thermo Scientific). Wash buffer contained PBS supplemented with 0.05% Tween20. Mean OD_450_ of blank wells for each antigen, which contained only coating antigen, were subtracted from raw OD_450_ values before analysis.

#### 2.6 Epitope mapping using an MSP2-derived peptide array

We constructed a custom PEPperCHIP® Peptide Microarray (PEPperPRINT) containing 128, 13-amino acid (13-mer) peptides present in a collection of 494 MSP2 variants sequenced previously using CCS (33). Amino acid sequences for MSP2 variants were digested (12 residue overlap) *in silico* (38) and the most frequent, distinct 13-mers were selected. Each of 16 subarrays contained poliovirus- (KEVPALTAVETGAT) and influenza haemagglutinin- derived (YPYDVPDYAG) peptides as positive controls. Sample plasmas were diluted 1:400. Unexposed plasma and subarrays containing only secondary antibody, were used as negative controls. Alexa Fluor™ 647-coupled Goat anti-Human IgG (H+L) Cross-Adsorbed Secondary Antibody (Invitrogen) was diluted 1:2000 for detection. Peptide microarrays were run according to manufacturer instructions. Briefly, slides were incubated with wash buffer (1x PBS, 0.05% Tween20) for 15 minutes and blocked (Rockland Blocking Buffer MB-070) for 30 minutes at room temperature. Plasma samples were diluted in staining buffer (1x PBS, 0.05% Tween20 and 10% Rockland Blocking Buffer) and incubated on the arrays overnight at 4°C. Slides were then washed 3 times before adding diluted detection antibody (1:2,000) for 45 minutes at room temperature. Slides were washed 3 times, dipped in buffer (1mM Tris buffer, pH 7.4) and air-dried. All incubation steps were done under constant shaking at 140 rpm. Slides were imaged using an Innopsys InnoScan 1100 AL 3-channel ultra-high-resolution scanner and analysed using GenepixPro 6.0. Features were aligned and extracted using GenePix. A feature was classified as positive if it was at least 30 pixels in diameter, had a fluorescence of at least 100, and if

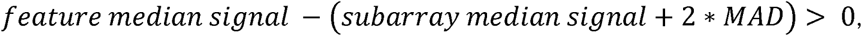

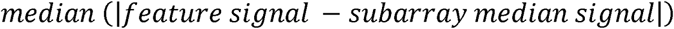

Both printed replicates of a feature within a subarray had to be positive for that feature to be considered plasma-bound or “recognized”. MSP2 peptides bound by negative controls were removed altogether from the analysis.

#### 2.7 Analysis and Statistical methods

OD_450_ values were batch-normalized, after subtracting blanks, using the following formula:

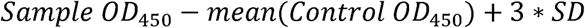

where *Control* is unexposed plasma pool, also called “negative control”. To identify plasmas binding to tested antigens, we applied a positivity cutoff using the mean negative control response of each antigen, in normalized OD_450_ plus 3 standard deviations. Any samples that did not pass the positivity cutoff for at least one of the ten antigens were removed from further analysis (N_Travelers_ = 0, N_Tanzania_ = 12). Breadth was defined as the number of variants bound after applying a cutoff of seropositivity. Breadth towards full MSP2 variants was determined separate from breadth towards truncated MSP2 variants. Normality of the data was checked by Shapiro-Wilk test and by examining the QQ plot on non-transformed data. Variance in normalized OD was determined by Wilcoxon test for pairwise comparisons. Heatmaps were plotted using pheatmap version 1.0.12 and the Complex Heatmap package for R (39); raincloud plots were created using ggdist for R (40,41). Figures were prepared using ggplot2 (42), patchwork (43) and ggpubr version 0.6.0 (44). Data analysis using R Studio was performed in R v 4.3.2. Alphafold2 (45) structures for the four MSP2 variants that were cloned and tested by ELISA were produced in Galaxy Version 2.3.1 (46). Oligomer structures for the same variants, as well as for the conserved termini of MSP2 were modelled in the Alphafold3 server (47). Signal peptides and GPI anchoring signals previously predicted with SignalP 6.0 (48) and big-PI Predictor (49) respectively, were removed from the amino acid sequences that were modelled. For structural modelling, serine and threonine residues were substituted by alanines in all N-Glycosylation sites predicted by NetNGlyc 1.0 (35).

### 3 Results

We assessed antibody responses of primary infected and previously exposed travelers as well as Tanzanian children and adults (**Figure S1)**, towards conserved and variable regions of MSP2 using IgG ELISA and peptide array. The four expressed MSP2 variants were isolated from travelers who recorded country/region of infection as Gambia (MSP2-FC27 1), South- East Asia (MSP2-IC 1), Ghana (MSP2-IC 2), and Kenya (MSP2-FC27 2) **(Table S2)**. In total, 98 Tanzanian samples were included, of which 49 were from a survey conducted in 1994 and 49 from a survey in 1999. Ages amongst the Tanzania participants ranged from 1 to 74 years old and 80 individuals (82%) were asymptomatic. Adult Travelers with acute febrile malaria, returning mainly from sub-Saharan Africa were also included (n=36) (**Table 1**). Travelers’ plasma samples for this study were collected 10 days after diagnosis and initiation of treatment, the timepoint previously shown to be the peak of IgG responses in this cohort (29).

**Table 1:**
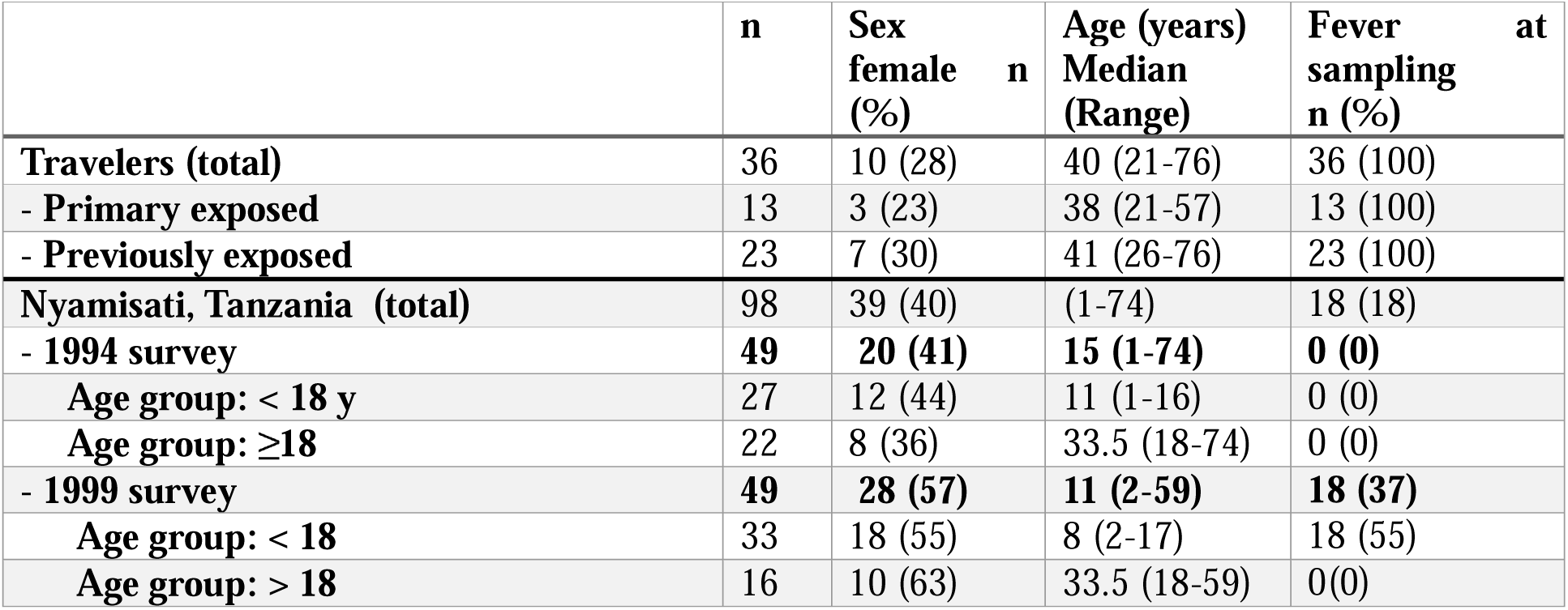
Characteristics of the study population.

#### 3.1 IgG responses and breadth of binding to full-length MSP2 variants increases after first exposure

We first measured the IgG antibody responses of both cohorts towards four recombinantly expressed full length MSP2 variants which were sequenced from patient isolates (**Table S2**). IgG antibody responses (normalized OD) followed a non-normal distribution as determined by QQ plot of residuals (**Figure S3A, B**), and by Shapiro-Wilk test for travelers (W = 0.602, p < 0.0001) and Tanzania samples (W = 0.691, p < 0.0001). IgG responses against full MSP2 antigens were significantly higher in previously exposed travelers than in primary infected travelers (p < 0.0001) (**Figure 1A**). Tanzanian samples were split into children (younger than 18 years) and adults (age 18 years or older). IgG responses to full MSP2 overall were not significantly different between age groups for 1994 samples (**Figure 1B**) while responses were significantly higher (p < 0.05) in adults compared to children for 1999 samples (**Figure 1C**). When comparing IgG responses towards specific MSP2 variants, we found no difference in responses to any of the four antigens for primary infected travelers (**Figure 1D**), while for previously exposed travelers there were higher responses to MSP2-FC27 2 than to MSP2-IC 2 (p < 0.05). For the 1994 survey in Tanzania, responses towards MSP2-FC27 2 were higher than those towards MSP2-FC27 1, MSP2-IC 1 and MSP2-IC 2 (p < 0.01, p < 0.001, p < 0.0001) (**Figure 1E**). For participants of the 1999 survey, responses showed a similar pattern, although responses to MSP2-IC 2 were significantly higher from those to MSP2-IC 1 (**Figure 1E**).

**Figure 1.**
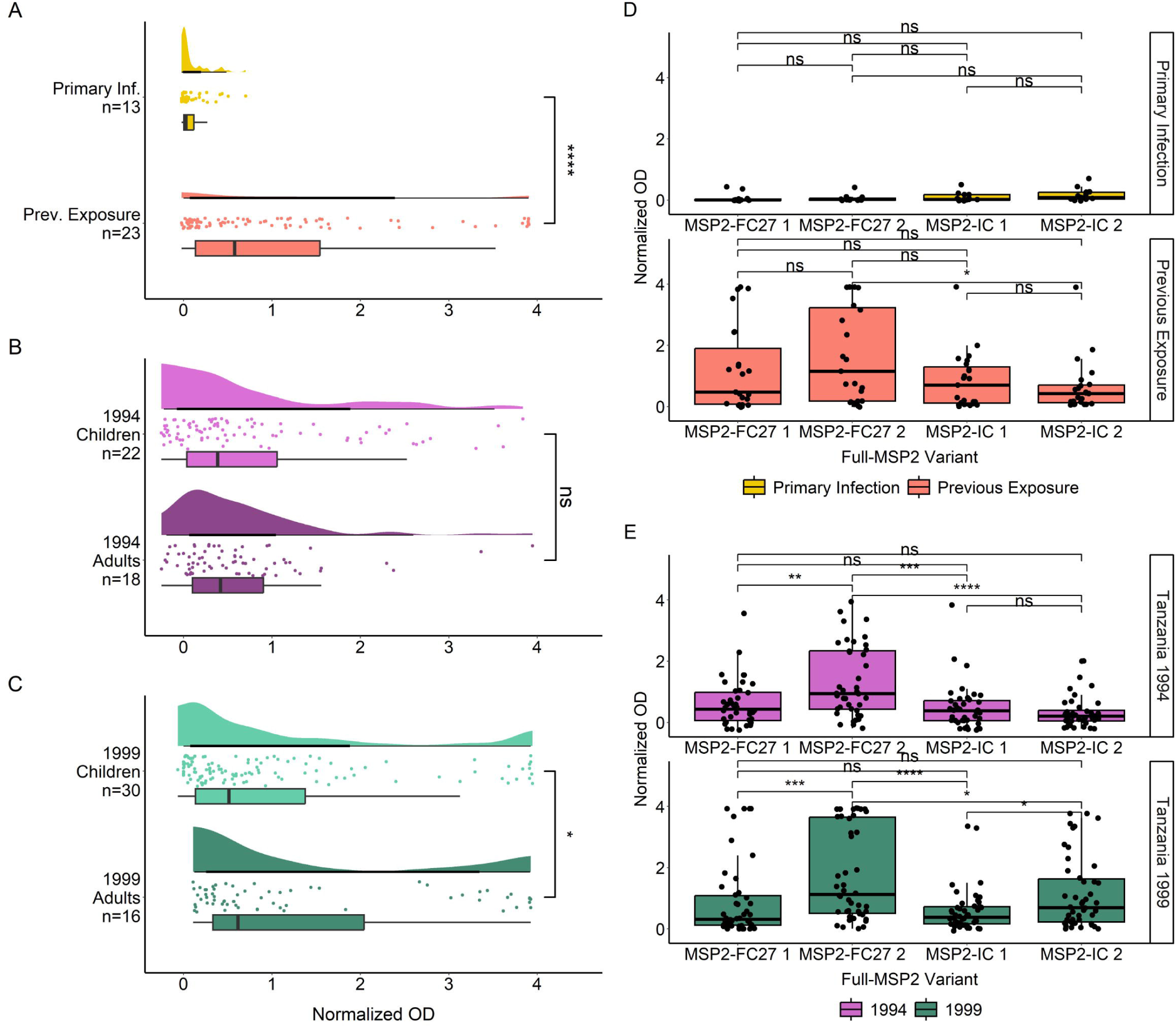
IgG responses towards full MSP2 variants by ELISA. Normalized absorbance of samples with detected IgG responses to at least one of the 10 tested antigens are shown in raincloud plots for Travelers (**A**), Tanzania 1994 (**B**) and Tanzania 1999 (**C**). Travelers are grouped by level of previous exposure to *P. falciparum* malaria and the Tanzania cohort is grouped by age. Boxplot lines denote median and CI .25-.75. IgG responses in normalized absorbance towards each of four MSP2 variants are shown for the same groups of travelers (**D**) and Tanzania samples (**E**). Normalized absorbance values were compared using unpaired Wilcoxon test where significance values are as follows: * < 0.05, ** < 0.01, *** < 0.001, **** < 0.0001.

Breadth towards full MSP2 variants was significantly higher in previously exposed travelers compared with primary infected travelers (p < 0.001) (**Figure 2A**). We were interested to see if individuals with higher breadth also had higher levels of anti-MSP2 IgG antibodies. We found that IgG responses in both primary infected and previously exposed travelers increased with greater breadth (**Figure 2B**). This was especially evident for previously exposed travelers with breadth of 4 compared with breadth of 3 or 2 full MSP2 variants (p < 0.0001, p < 0.01). We saw no difference in breadth between the Tanzanian subgroups—1994 children and adults, and 1999 children and adults (**Figure 2C**). However, we did see increasing normalized OD associated with greater breadth for all four Tanzanian subgroups (**Figure 2D**), similar to the pattern observed for travelers. Tanzania 1994 children and adults had similar IgG levels with breadth 3 and 4, whereas Tanzania 1999 children and adults responded significantly higher when they displayed breadth of 4 (p < 0.01).

**Figure 2.**
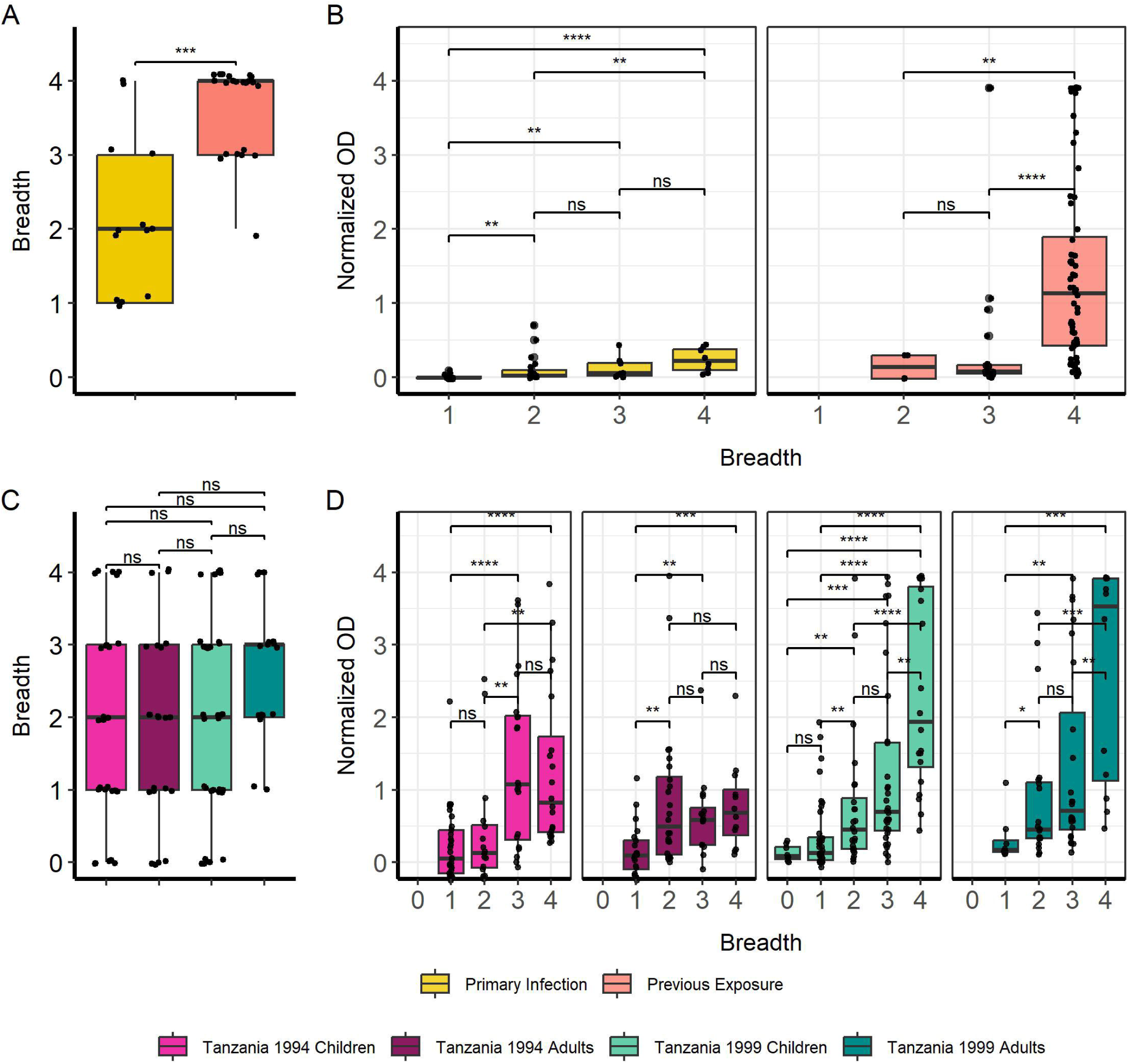
Increasing breadth towards full MSP2 variants is associated with increased IgG responses. Breadth for each individual sample was plotted, where samples were considered “binding” if they passed the positivity cutoff based on negative controls for travelers (**A**) and Tanzania samples (**C**). Boxplots denote CI 0.95 and mean breadth. IgG responses were plotted against breadth to full MSP2 variants for travelers (**B**) and Tanzania samples (**D**). Pairwise comparisons performed with Wilcoxon test. Significant differences are marked with asterisks (* < 0.05, ** < 0.01, *** < 0.001, **** < 0.0001).

#### 3.2 Levels of IgG antibodies and breadth of binding to truncated MSP2 variants increases with first exposure

We then evaluated the binding of naturally acquired IgG antibodies to truncated MSP2 variants lacking their conserved termini (**Figure 3**). Similar to what we observed for the full variants, we found significantly higher IgG responses in previously exposed compared to primary infected travelers (p < 0.0001) (**Figure 3A**). Furthermore, Tanzanian adults had overall similar IgG responses compared with children for the 1994 survey (**Figure 3B**) but overall higher IgG responses compared with children for the 1999 survey (p < 0.05) (**Figure 3C**). When comparing responses to truncated MSP2 variants, IgG responses in primary infected travelers were very low, although responses were significantly higher towards MSP2- IC 1 than MSP2-FC27 1 (p < 0.01) and MSP2-FC27 2 (p < 0.001) (**Figure 3D**). Responses towards MSP2-IC 2 were also higher than those towards MSP2-FC27 1 (p < 0.05) and MSP2- FC27 2 (p < 0.01). Plasma IgG from previously exposed travelers, interestingly, responded significantly higher to MSP2-IC 1 than MSP2-FC27 1 but no other significant comparisons were found (**Figure 3D**).Tanzania 1994 responses were significantly higher towards truncated MSP2-FC27 2 than towards MSP2-FC27 1 (p < 0.0001) and MSP2-IC 2 (p < 0.01); higher towards MSP2-IC 1 than towards MSP2-FC27 1 (p < 0.0001) and MSP2-IC 2 (p < 0.001); and no significant difference was observed between MSP2-FC27 2 and MSP2-IC 1 (**Figure 3E**). Tanzania 1999 IgG responses towards MSP2-FC27 2 were higher than towards MSP2- FC27 1 (p < 0.001); IgG responses towards MSP2-IC 1 were significantly higher than towards MSP2-FC27 1 (p < 0.0001) and MSP2-IC 2 (p < 0.05). Additionally, responses towards MSP2-IC 2 were higher than responses to MSP2-FC27 1 (p < 0.01) and responses to MSP2-IC 1 were higher than responses to MSP2-IC 2 (p < 0.05) (**Figure 3E**).

**Figure 3.**
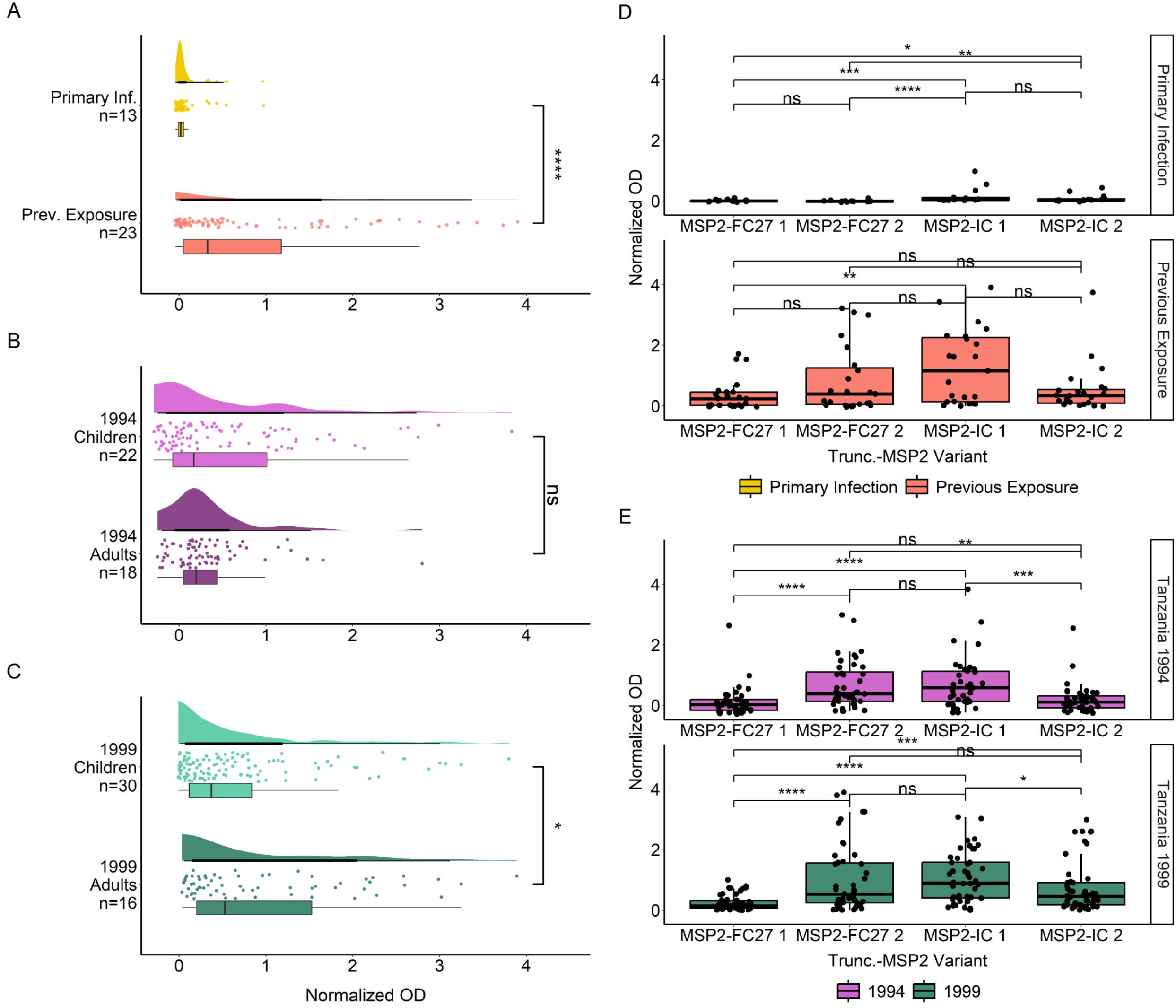
IgG responses towards truncated versions of MSP2 antigens. IgG responses towards truncated MSP2 variants by ELISA. IgG responses in normalized absorbance are shown for travelers (**A**), as well as Tanzania participants in the 1994 (**B**) and 1999 (**C**) surveys, respectively. Traveler samples are grouped by level of previous exposure to *P. falciparum* malaria while Tanzania samples are grouped by age. IgG responses in normalized absorbance towards each of four MSP2 variants are shown for the same groups of travelers (**D**) and Tanzania samples (**E**). Normalized absorbance values were compared using unpaired Wilcoxon test where significance is marked as follows: * < 0.05, ** < 0.01, *** < 0.001, ****< 0.0001.

Breadth of IgG responses towards truncated MSP2 variants was again significantly higher in previously exposed compared to primary infected travelers (p < 0.001) (**Figure 4A**). IgG responses towards truncated MSP2 variants also increased with greater breadth for previously exposed individuals, while no comparisons could be made for primary infected individuals due to the low number of observations (**Figure 4B**). Unlike breadth comparisons for full MSP2, we found a significantly higher breadth towards truncated MSP2 variants in adults from the 1999 survey compared with adults from the 1994 survey (p < 0.01) and children from the 1994 survey (p < 0.05) (**Figure 4C**). There was no difference in breadth to truncated MSP2 between children and adults from the 1994 survey, nor between children and adults from the 1999 survey (**Figure 4C**). Finally, IgG responses were significantly increased in Tanzanian individuals who had greater breadth to truncated MSP2 variants (**Figure 4D**), similar to the pattern observed for IgG responses towards full MSP2 variants.

**Figure 4.**
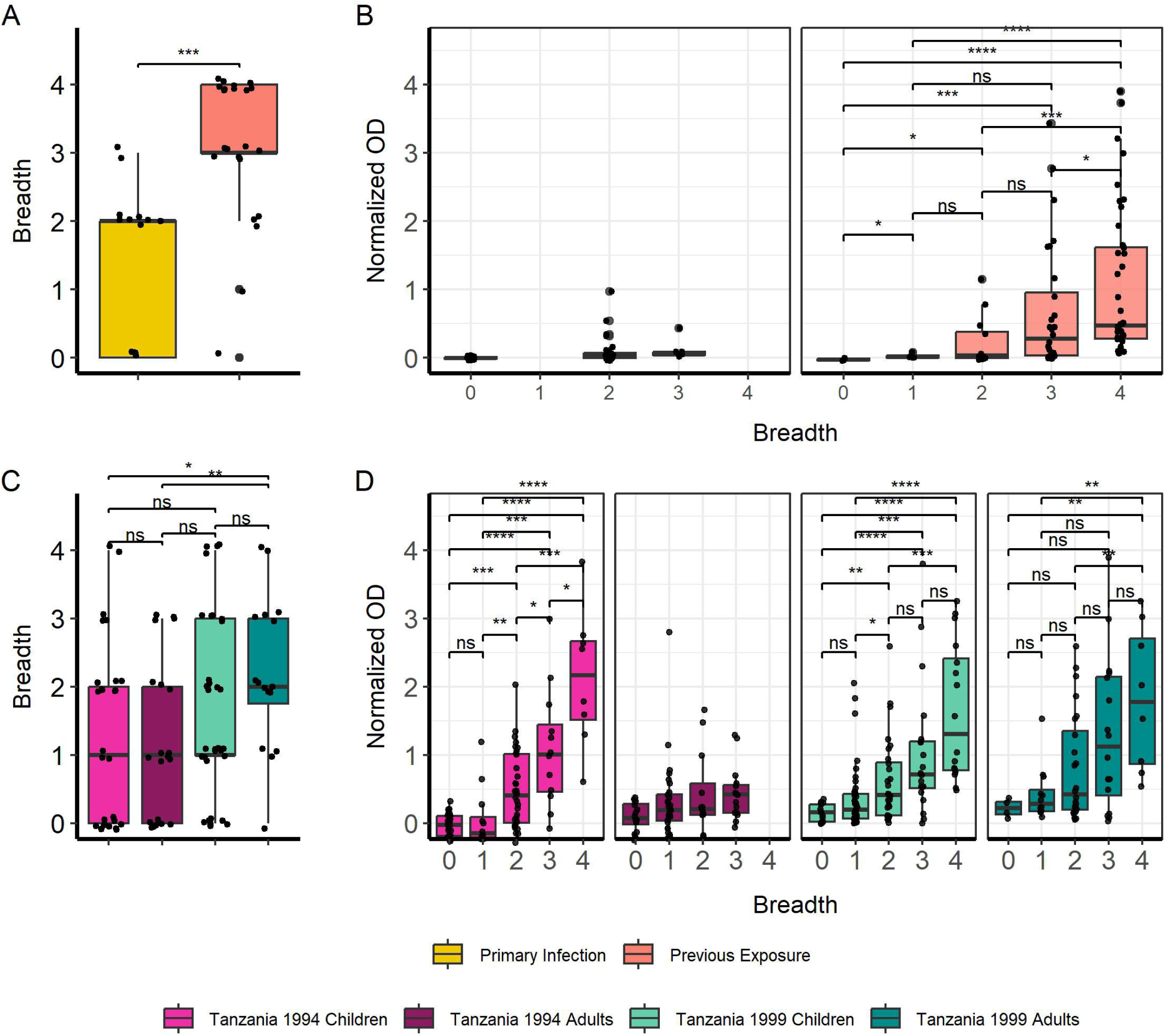
Increasing breadth towards truncated MSP2 variants is associated with increased IgG responses. Breadth for each individual was plotted, where samples were considered “binding” if they passed the positivity cutoff based on negative controls for travelers (**A**) and Tanzania samples (**B**). Boxplots denote 0.95 CI and mean breadth. IgG responses in normalized absorbance to all four truncated variants were plotted against increasing breadth to truncated MSP2 variants for travelers (**C**) and Tanzania samples (**D**). Pairwise comparisons performed with Wilcoxon test and significance is show with asterisks (* < 0.05, ** < 0.01, *** < 0.001, **** < 0.0001).

#### 3.3 IgG antibodies that recognize conserved MSP2 terminus peptides are not a substantial part of the humoral response in malaria

To investigate the binding of IgG antibodies to the conserved termini of MSP2 alone, we tested the plasma samples against two peptides covering the complete conserved N- and C-termini of the native antigen (**Figure 5**). There were no significant differences in IgG responses towards the peptides when comparing primary infected and previously exposed travelers (**Figure 5A**). There was a significantly higher response in Tanzania 1994 adults compared with children (p < 0.05) (**Figure 5B**). There were no significant differences in IgG binding towards the conserved peptides when comparing adults and children from the Tanzania 1999 survey (**Figure 5C**). In all three groups the absorbances measured were negligible. IgG responses towards the conserved N terminus were significantly higher than those directed towards the C terminus in both primary infected and previously exposed travelers (both p < 0.001) although negligible (**Figure 5D**). No such significant difference was found for Tanzania 1994 (**Figure 5E**) or Tanzania 1999 individuals (**Figure 5E**).

**Figure 5.**
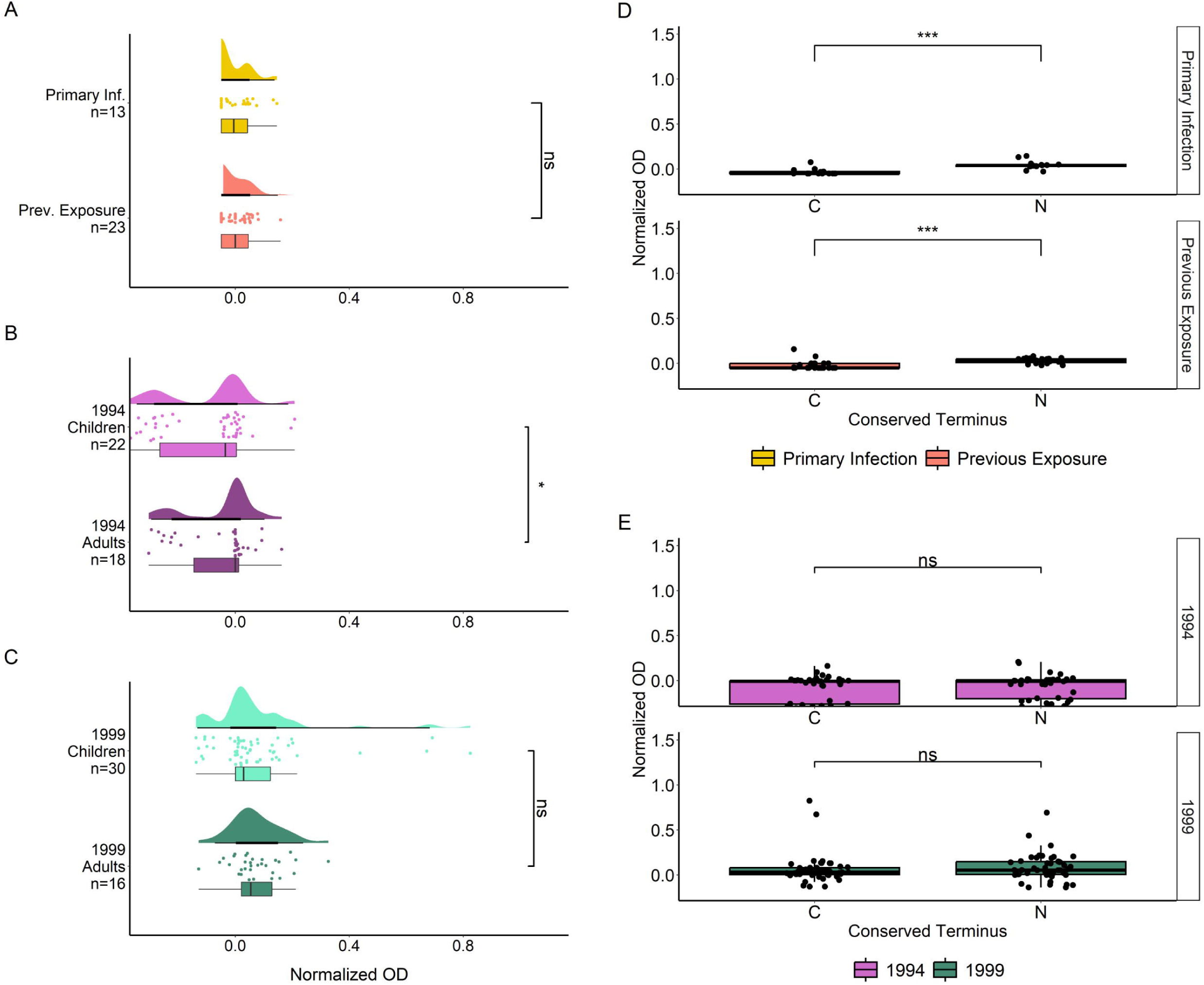
IgG responses towards conserved MSP2 termini by ELISA. IgG responses to N and C terminal peptides, in normalized absorbance, are shown for travelers (**A**), Tanzania 1994 (**B**) and Tanzania 1999 (**C**), where travelers samples are grouped by level of previous exposure to *P. falciparum* malaria and Tanzania samples are grouped by age. IgG responses in normalized absorbance towards both peptides are shown for the same groups of travelers (**D**) and Tanzania samples (**E**). Normalized absorbance values were compared using unpaired Wilcoxon Test where significance values are as follows: * < 0.05, ** < 0.01, *** < 0.001, **** < 0.0001.

We also evaluated the binding of IgG in plasmas from the four travelers who were infected with *P. falciparum* parasites from which the respective four MSP2 variants were sequenced and expressed **(Figure S4)**. Traveler 10 (MSP2-FC27 1), traveler 3 (MSP2-FC27 2), traveler 46 (MSP2-IC 1), and traveler 35 (MSP2-IC 2) recognized the full version of their infecting MSP2 variant; IgG from traveler 35 (MSP2-IC 2) and traveler 46 (MSP2-IC 1) could also recognize full MSP2-FC27 2 **(Figure S4A)**. Traveler 35 recognized the truncated version of MSP2-FC27 2 and MSP2-IC 1 but not their own infecting MSP2 variant (MSP2-IC 2), indicating a level of cross-reactivity to the two allelic families. Traveler 46 recognized truncated MSP2-FC27 2 and the truncated version of their own variant (MSP2-IC 1). CCS of Traveler 46 showed the presence of both IC and FC27 clones in the isolate, explaining the recognition of both MSP2 families by this subject’s plasma **(Figure S4B)**. Traveler 10 and traveler 3, both harboring FC27-type MSP2, were unable to recognize any truncated MSP2 variants, suggesting IgG binding to conserved epitopes. Interestingly, none of the tested travelers recognized N- or C-terminus peptides by ELISA, and all four responded at the same level as evidenced by overlapping points **(Figure S4C)**.

A binary response matrix was created to visualize all antigens bound by individual plasmas (**Figure S5)**, including those who did not respond to any antigens (N=9 _Tanzania1994_, N=3 _Tanzania1999_), according to the seropositivity cutoff values for each group. Two of the 13 tested primary infected travelers recognized the N-terminus peptide, and 2 of the 23 tested previously exposed travelers could bind the C-terminus (**Figure S5A**). Only one child from the Tanzania 1994 survey was able to recognize the conserved N- terminus (**Figure S5B**). From the 1999 survey, IgG from one child (p67) recognized both the N- and C- terminus, while one child (p118) and one adult (p106) recognized the N-terminus only (**Figure S5C**). We also visualized paired IgG responses towards full and truncated versions of individual MSP2 variants for single individuals (**Figure 6**). Primary infected travelers show significantly higher responses to full MSP2-FC27 2 compared with truncated MSP2-FC27 2 (p < 0.01), with no other significant differences being observed **(Figure 6A)**. Previously exposed travelers showed higher IgG responses to full MSP2-FC27 1 and MSP2-FC27 2 compared with the truncated versions (both p < 0.05). No significant difference was found for full and truncated versions of the two IC variants **(Figure 6A)**. Tanzanian children and adults from the 1994 survey showed significantly higher responses to full MSP2-FC27 1 compared with truncated MSP2-FC27 1 (p < 0.01, p < 0.05). Other comparisons were not found to be significant **(Figure 6B)**. Tanzanian children from the 1999 survey, showed significantly higher responses to full MSP2-FC27 2 (p < 0.05) and significantly lower responses to full MSP2-IC 1 (p < 0.05), than their truncated counterparts. Adults from the same survey had significantly higher responses to full MSP2-FC27 1 than the truncated version (p < 0.01). Furthermore, adults also showed significantly higher responses to truncated MSP2-IC 2 than full MSP2-IC 2 **(Figure 6C)**.

**Figure 6.**
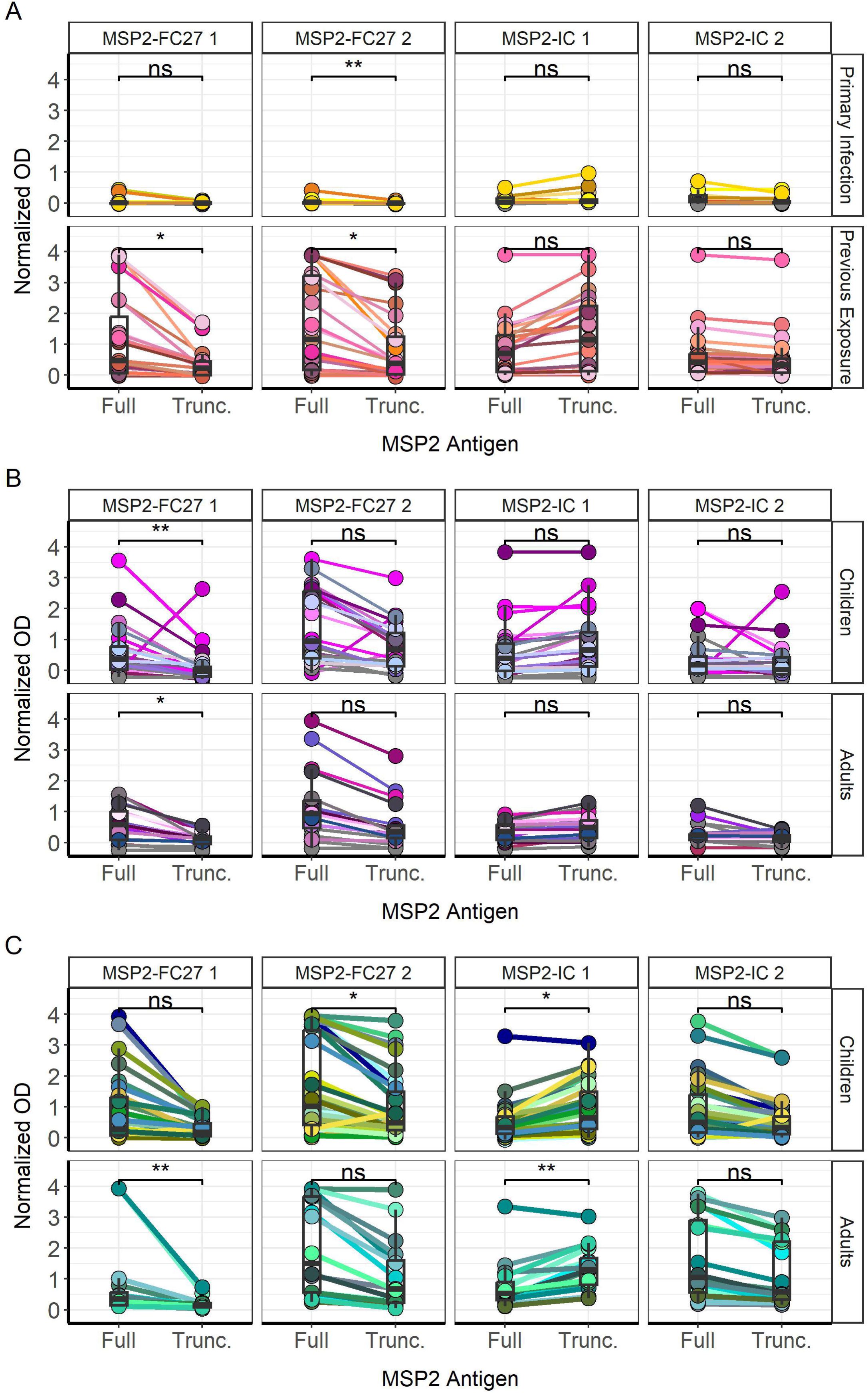
Differences between IgG responses towards full and truncated MSP2 variants. IgG responses for each sample, indicated with connecting lines, were plotted against each antigen tested for primary infected and previously exposed travelers (**A**). Responses were also plotted for children and adults in the 1994 (**B**) and 1999 (**C**) surveys in Tanzania. Comparisons of mean normalized absorbance were performed by unpaired Wilcoxon test (* < 0.05, ** < 0.01, *** < 0.001, **** < 0.0001).

We then plotted IgG responses to full MSP2 variants, in normalized OD, against IgG responses to truncated MSP2 variants and performed a Spearman rank-order correlation for each study group (**Figure S6**). A significant, positive correlation was found for primary infected and previously exposed travelers (**Figure S6A**) as well as for children and adults from Tanzania 1994 (**Figure S6B**) and 1999 surveys (**Figure S6C**). Given that the Tanzania 1999 children and adults showed significant differences in median IgG response towards full and truncated MSP2 overall, we performed a Spearman rank-order correlation to analyze anti- MSP2 IgG levels (normalized OD) with increasing age (**Figure S7**). The relationship between IgG responses towards full or truncated MSP2 variants, and the age of individuals from Tanzania 1994 and 1999, was variant specific and dependent on age group. (**Figure S7A-D**). We found no significant correlation between IgG response and age for the conserved termini for Tanzania 1994 (**Figure S7E**) or 1999 (**Figure S7F**).

We also stratified IgG responses in Tanzania study participants by fever and microscopy positivity status (**Figure S8**). We found no significant differences in IgG responses towards full MSP2 variants for children who were microscopy negative or positive (**Figure S8A**). This was also the case for truncated MSP2 variants (**Figure S8B**) and conserved MSP2 termini (**Figure S8C**). Similarly, we found no significant differences in IgG responses towards full (**Figure S8D**) or truncated (**Figure S8E**) MSP2 variants or the termini (**Figure S8F**) for adults who were microscopy positive or negative at sampling. No Tanzanian adults in either survey had a recorded fever at the time of sampling. We only found significant differences in IgG responses between children who were febrile and non-febrile at the time of sampling for full MSP2-IC 2 (**Figure S8G**) and for truncated MSP2-FC27 1 (**Figure S8H**). We did observe a significantly greater (p < 0.01) response to both N- and C- terminus peptides in Tanzanian children who were febrile at sampling, compared with those who were nonfebrile (**Figure S8I**).

#### 3.4 Binding to linear 13mer epitopes from the conserved termini of MSP2 in a selection of travelers

We evaluated the binding of IgG antibodies to conserved 13 amino acid linear peptides from MSP2 by peptide microarray (schematic, **Figure S9**). Four primary infected travelers, all tested by IgG ELISA in this study, were selected for analysis; all were Swedish individuals without prior malaria and without treatment failure. Array slides were printed with the 128 most frequent 13-mer peptide sequences (single amino acid overlap) present in a collection of 494 MSP2 variants that were previously sequenced for a separate study. We defined bound epitopes with single-residue resolution using the number of overlapping peptides bound by the tested plasma (coverage) ranging from 0 to 8 (**Figure 7**). Traveler 6 IgG bound the N terminus amino acids “YSNTFINNAYN” in 6-8 overlapping linear peptides, while traveler 20 IgG bound the N terminus amino acids “YSNTFINNA” in 3-5 overlapping linear peptides (**Figure 7A**). Traveler 22 IgG bound the N terminus amino acids “TFINNAY” with low overlap (2–3) while traveler 35 did not bind any peptide derived from the N terminus (**Figure 7A**). Only IgG from traveler 22 and traveler 35 recognized full MSP2-IC 1 (traveler 22) and full MSP2-IC 2 (traveler 22 and traveler 35) by ELISA (**Figure 7B**). Traveler 22 recognized the conserved N-terminus by ELISA. The location of plasma IgG binding and amino acid overlap are shown on AlphaFold2 models of the full antigens used in IgG ELISA (**Figure 7C**). Finally, we observed that each of the four plasmas displayed unique patterns of epitope binding.

**Figure 7.**
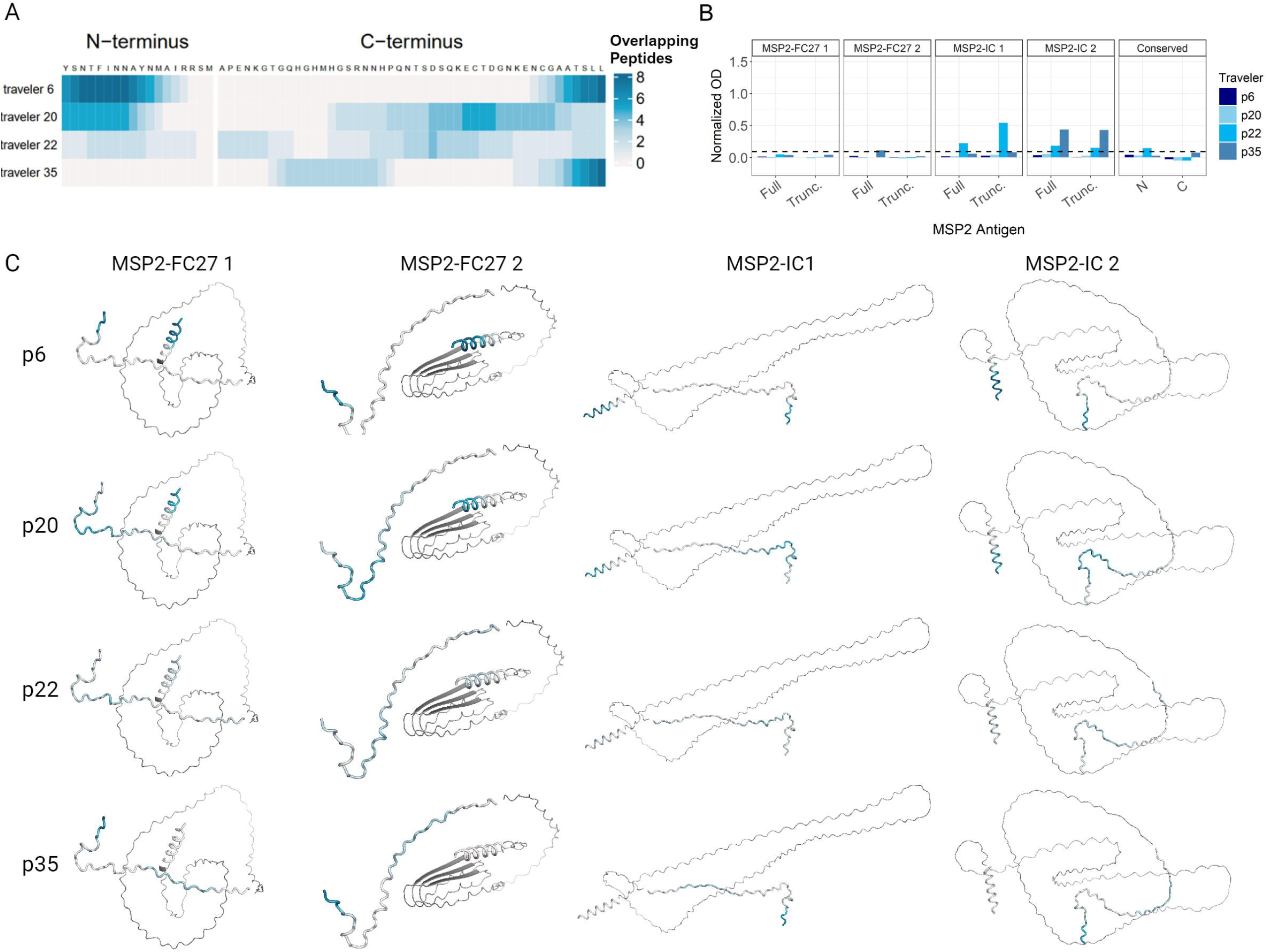
Binding to N- and C-terminus 13mer peptides by four primary infected Travelers. Number of positive, overlapping 13-mer peptides were mapped to each amino acid residue of the N- and C- termini on the peptide array for each of four primary infected travelers (**A**). IgG responses in normalized absorbance for each traveler against all ten MSP2 antigens (**B**). Dashed line indicates global mean negative cutoff. Positive overlapping 13mer peptides for each amino acid of the N- and C-termini for each traveler mapped onto AlphaFold2 predictions of the full MSP2-FC27 4, MSP2-FC27 1, MSP2-IC 1, and MSP2-IC 2 antigens used in the ELISA assays (**C**).

#### 3.5 Modelling of MSP2 variants using AlphaFold3 predicts structural heterogeneity

The subtle differences observed in the recognition of the conserved termini by IgG in plasma from travelers and Tanzanian individuals, when tested as separate peptides or on the native antigen, suggested context-dependent differences in the 3D structure of these regions. SDS-PAGE under denaturing conditions showed that the purified recombinant antigen produces disulfide bond-independent, heat- and detergent-stable dimers or trimers that might affect the structure or the accessibility of epitopes in the conserved termini (**Figure 8**). Though we cannot assert the oligomerization capacity for the N terminus peptide by SDS-PAGE, we observed that the C terminus ran at approximately 15 kDa, consistent with the formation of heat- and detergent-stable trimers predicted to be 14.7 kDa in size (**Figure 8B**).

**Figure 8.**
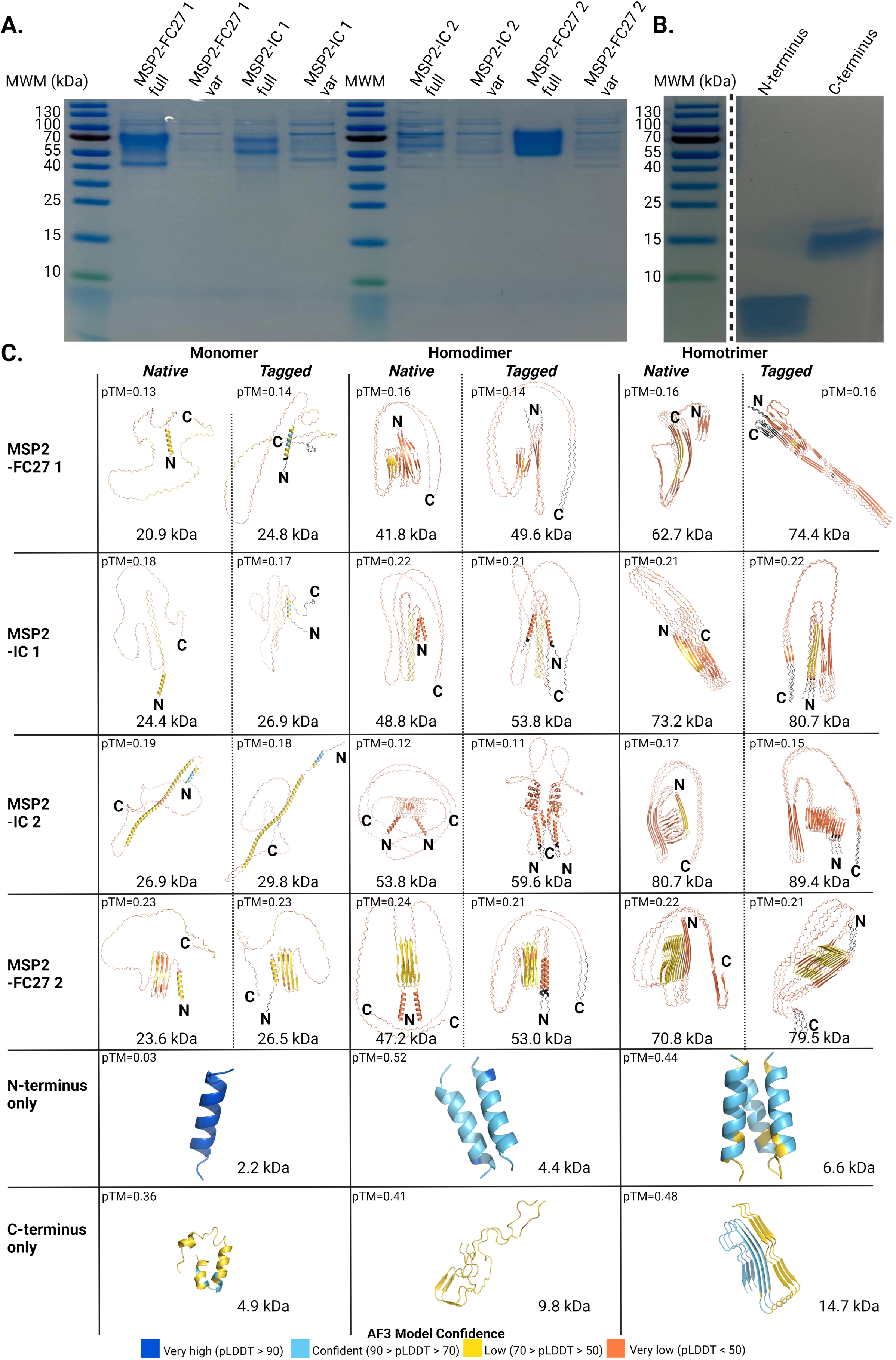
Structural heterogeneity in the conserved termini of MSP2 is associated to its oligomerization status. SDS-PAGE for (**A**) recombinant proteins and (**B**) synthetic peptides used in the characterization of antibody responses against MSP2 in the presence or absence of the reducing agent DTT, respectively. (**C**) Alphafold3 models for monomers and oligomers of the native and engineered MSP2 variants used in the study. AF3 models for monomeric and oligomeric forms of the synthetic peptides corresponding to the conserved N and C termini are also presented. Predicted molecular weight and pTM scores are shown for every model. Truncated MSP2 variants are marked as “var”. Tags engineered into the variants before expression are shown in black. Colors represent pLDDT scores calculated for every residue in the model.

We then modelled the structure of monomers, dimers and trimers of the native and tagged- engineered versions of the four full constructs with Alphafold3 (AF3), as well as the structure of monomers, dimers and trimers formed by the conserved termini alone (58). Predicted local distance difference test (pLDDT) scores for most residues in structured regions containing alpha-helices or beta strands ranged between 50 and 70. All monomers for the recombinant variants used in this study showed conserved N termini with alpha-helical and disordered conformations (**Figure 8C**). Likewise, monomers and oligomers of the conserved N terminus alone were predicted to assume an alpha-helical fold (**Figure 8C**). In addition, the N terminus of homodimer subunits formed by MSP2-IC 1, MSP2-IC 2 and MSP2-FC27 2 was shown to be alpha helical. In contrast, the conserved N terminus domains of all the computed homotrimers and the homodimer formed by MSP2-FC27 1 were found to form beta sheets including one beta strand from every subunit in the complex.

Finally, the C-terminus of all monomers and homodimers showed an intrinsically disordered structure. On the contrary, the C termini in homotrimers assumed a beta sheet conformation for the two FC27 variants tested while keeping a disordered conformation in MSP2-IC 1 and MSP2-IC 2 (**Figure 8C**). Engineering tags into the expression constructs only seems to affect the predicted conformation of the conserved C terminus in homotrimers formed by variants MSP2-IC 1 and MSP2-FC27 2, where the tagged version of these variants assumes a beta sheet conformation in contrast to the disordered fold observed in untagged homotrimers. Structure predictions for monomers and oligomers of the C-terminus domain alone display a high degree of structural heterogeneity, with monomers formed predominantly by alpha helices (59.6% of the peptide or 28 residues out of 47) and a smaller fraction of disordered residues (40.4%); and dimers being 85.1% disordered (40 residues out of 47 in every subunit) and 14.9% beta strands (7 residues) (**Figure 8C**). Structural prediction of homotrimers for the conserved C terminus only (the stable 15 kDa complex observed by SDS-PAGE, **Figure 8B**), revealed a dominant beta sheet component formed by four beta strands per subunit including 25 out of the 47 residues (53.2%) (**Figure 8C**).

### 4 Discussion

A more detailed knowledge of the immune response towards polymorphic *P. falciparum* antigens is needed to understand naturally acquired immunity to malaria, and to improve vaccine design. MSP2 has been studied for several decades as a potential vaccine candidate (50), but its extensive polymorphism has remained a major challenge. Reference lab isolates of *P. falciparum* such as NF54 and 3D7 have been used historically to measure MSP2-specific antibody responses, but they do not accurately represent the diversity of MSP2 found in field isolates (33). New high fidelity sequencing methods allowed us to accurately sequence and recombinantly express MSP2 variants from *P. falciparum* isolates, to better understand naturally acquired IgG responses towards MSP2 from the infecting parasite.

We did not find substantial levels of naturally induced antibodies against the N- or C-terminus of MSP2 variants sequenced from parasite isolates and expressed as peptides. We found increased IgG responses and breadth in plasma from previously exposed travelers compared with primary infected travelers, towards both full and truncated MSP2 antigens. Previous studies also showed increased IgG responses towards *P. falciparum* antigens in previously exposed compared with primary infected travelers (32), and specifically for MSP2 from reference strain FC27 (51). Tanzanian children and adults in the 1994 survey had similar levels of IgG responses to full and truncated MSP2 antigens. This was unexpected since antibody responses have been shown to increase with repeated exposure (52); along with breadth, antibody titers also increase as a function of exposure to pathogens using antigen variability as immune escape mechanisms (37,53). In comparison, we did find greater IgG responses towards full and truncated MSP2 in adults compared with children from the Tanzania 1999 survey. The most likely explanation for this difference between surveys is the age cutoff (18 years) used for adults and children; In highly endemic areas, clinical immunity typically occurs at a younger age (54,55), so including older children and teenagers in the “children” group likely masked the effect of age. Median breadth towards full and truncated MSP2 was also not significantly different between age groups. Greater breadth was, however, associated with increased IgG responses, for both full and truncated MSP2 variants and for both travelers and Tanzanian individuals, indicating a relationship between breadth of response and intensity of response towards MSP2 variants regardless of exposure level. Importantly, we rely on self-reported previous malaria infection for travellers; therefore, it is impossible to know if those with higher breadth are also those who experienced a higher number of infections.

Very few individuals tested were able to recognize the conserved N- and C- terminus peptides by ELISA. It is important to note that the median IgG, in normalized OD, was negligible for travelers in addition to both Tanzania surveys, indicating that while few individuals may be able to recognize the conserved regions, they are not representative of the anti-MSP2 humoral responses in the population. The high number of asymptomatic infections in the cohort and the high prevalence of *P. falciparum* at the time of survey, coupled with our observation of limited IgG targeting the conserved regions, support the idea that conserved MSP2 regions are not important for humoral immunity. Previous studies have shown that mouse monoclonal antibodies generated by immunization with recombinant MSP2 recognized epitopes in the conserved termini of the antigen (23). Repeats were also shown to be antibody-bound in mice (56). A 3D7-based vaccine trial demonstrated strain-specific recognition of the repetitive and dimorphic regions, whereas the conserved domains of the antigen were not immunogenic (57). A more recent phase 1 clinical trial of a vaccine including two allelic forms of MSP2 showed that immunized participants had antibodies targeting the C-terminus regions of 3D7 and FC27 (58). Further studies will be needed in order to assess the differences in the recognition patterns of vaccine-induced (previous studies) and naturally-acquired (reported here) antibodies against MSP2. IgG responses to the conserved termini may be reliant on epitope accessibility and protein structure (59); they may also be rare due to structural heterogeneity or low affinity (60).

If anti-MSP2 responses depended on recognition of family-specific dimorphic sequences, we would expect an individual harboring a variant of one family to show cross-reactivity towards other variants of that family. This was not that case for our autologous traveler plasmas. Traveler 10, infected with a parasite expressing MSP2-FC27 1, could not recognize MSP2- FC27 2. Likewise, traveler 46, infected with a parasite expressing MSP2-IC 1, could not recognize MSP2-IC 2. Variant-specific responses suggest humoral immunity against MSP2 depends on the recognition of variable region epitopes. This would translate into the need for individuals to be exposed to many different variable regions, or MSP2 variants, to successfully target MSP2. A previous study evaluating plasmas collected soon after diagnosis, however, showed that naturally acquired antibodies recognize recombinant polymorphic domains, with some degree of intra-family cross-reactivity for IC but no intra-family cross-reactivity for FC27 (61). Differences in the timing for sample collection might explain conflicting results for the IC family, with earlier antibody responses (plasmas collected soon after diagnosis) targeting relatively conserved dimorphic regions of the antigen and IgG at 10 days post-diagnosis (this study) recognizing the more polymorphic central domain.

The cross-sectional Tanzanian surveys limit us from drawing connections between previous episodes and IgG responses. A previous study on the Tanzanian cohort showed that levels of antibodies against multiple *P. falciparum* antigens were higher in 1999 than 2010, a lower transmission period (62). We expected to find a similar trend of higher IgG responses in 1994 compared to 1999 but did not. Since both surveys occurred during high transmission years, the change in prevalence may not have significantly affected antibody levels in the population. We also did not find a significant correlation of increasing anti-MSP2 IgG with age, which contradicts results from the same cohort in 1999 and 2010, where responses of children age 0 to 15 against MSP2 from strains Dd2 and CH150/9 were evaluated (62), but matches results from a study showing that in a hypo endemic, continuously exposed population in Brazil, there was no clear age-dependent increase in prevalence or mean levels of specific IgG (56).

MSP2 is largely intrinsically disordered and IgG binding to MSP2 occurs largely via linear epitopes, but previous work suggests that the structure of the antigen could be different when anchored by the C-terminus to a lipid membrane, thus modulating the presentation of N-terminus epitopes to the immune system (63). Detailed structural analysis has also demonstrated significant conformational differences between recombinant and soluble MSP2, and the antigen expressed on the merozoite surface, which could affect the efficacy of a vaccine based on recombinantly produced MSP2 (23). Immunization of mice with recombinant MSP2 has been shown to induce antibodies against the conserved termini which are unable to bind to the native protein as present on the parasite’s surface (64). Contradictory results showed that mouse antibodies cross-recognized native MSP2, in a family specific manner, after immunization with recombinant peptides (56). The plasmas used in our study contain antibodies elicited by natural infection where individuals are exposed to MSP2 in its native, membrane-anchored, conformation and, possibly also, to antigen fragments shredded during the lysis of infected erythrocytes and the clearance of the parasite by the immune system. Recognition of recombinant, soluble MSP2 by these individuals indicates that conformational differences may not inhibit all IgG binding of linear epitopes.

While MSP2-IC 1 and MSP2-IC 2 are relatively similar antigens (**Table 1**), they differ in the number of repeats: MSP2-IC 1 contains 9 repeats of the motif “SAGG” while MSP2-IC 2 contains 26. We found higher IgG responses towards full and truncated MSP2-IC 1 compared with full and truncated MSP2-IC 2 for Tanzania 1994 and Tanzania 1999. The number of “SAGG” repeats was previously shown to be important for the recognition of variants of the IC family by mouse monoclonal antibodies (12), while similar studies also showed the importance not only of the presence but also the number of FC27 family repeats (65). While testing full length MSP2 variants by ELISA, we observed that previously exposed travelers, Tanzania 1994 and Tanzania 1999 individuals recognized MSP2-FC27 2 more than the other 3 variants. Responses to MSP2-FC27 1 and MSP2-IC 1 were similar in all 3 groups as well. While testing truncated MSP2 variants, we observed that previously exposed travelers recognized MSP2-IC 1 and MSP2-FC27 2 at the same level. This was also true for Tanzania 1994 and 1999 individuals. IgG responses towards MSP2-FC27 1 and MSP2-IC 2 were lower in all 3 groups, and similar for previously exposed travelers and Tanzania 1994.

When grouped and colored by individual samples, it was clear that several individuals from all 4 exposure groups were able to recognize both full and truncated versions of the same tested antigen, at very similar levels. In addition, IgG to truncated MSP2-IC 1 was greater than to full MSP2-IC 1 in the 1999 survey from Tanzania, although no significant differences were observed for previously exposed travelers and Tanzania 1994. The observation that one MSP2 variant from each allelic family (MSP2-FC27 2 and MSP2-IC 1) was more recognized by highly exposed individuals, suggests that dimorphic sequences are not wholly responsible for anti-MSP2 humoral immunity. Overall, results suggest that IgG responses to MSP2 are dependent on recognition of the variable region and are not limited to dimorphic parts of the antigen. The number of repeats, and thus protein length, is likely important for structural conformation, structural heterogeneity, and the ability of humoral responses to recognize protective epitopes.

The low predicted template modelling (pTM) scores obtained reflect both the ability of AF3 to predict protein disorder and its limitations in the prediction of a precise topology for the predicted disordered regions (66). Structural modelling with AF3 showed a wide range of context-dependent structural heterogeneity in the conserved termini of MSP2 and the full antigen. This heterogeneity might explain the differences in the magnitude of IgG responses towards the conserved domains when in the context of the native, possibly oligomeric, recombinant antigen or when presented as oligomeric synthetic peptides in the absence of the central, polymorphic and intrinsically disordered domains. Though antibody clones capable of binding some conformations of the conserved termini might be present in the plasma, these don’t seem to bind the intrinsically disordered or beta sheet-rich structures predicted in the recombinant oligomers tested by ELISA in this study. Epitope masking has been proposed as a mechanism that prevents antibodies from recognizing conserved, strain-transcending epitopes on the termini domains of membrane-bound MSP2 (67). 6D8 is a mouse monoclonal antibody produced by immunization with oligomeric MSP2 (64) that binds the N-terminus of the recombinant soluble protein but that fails to recognize the membrane-bound form of the antigen (67). In this study, we present evidence demonstrating the formation of heat- and detergent-stable oligomers for MSP2-derived recombinant proteins and peptides in the presence or absence of the reducing agent DTT. Oligomerization in the presence of DTT indicates that the formation of disulfide bridges between cysteines in the conserved termini of separate MSP2 subunits does not play a role in the generation of the antigen dimers or trimers observed. Furthermore, we show that oligomerization, especially the formation of homotrimers, induces structural changes in the conformation of the N terminus. 6D8’s inability to bind MSP2 on the merozoite surface suggests that this antigen could be found as a monomer *in vivo*, with the N terminus assuming an alpha helical conformation. This could explain the minimal, yet significant, binding observed for plasmas from exposed individuals on the synthetic peptide covering the protein’s N terminus, that regardless of its oligomerization status, retains an alpha helical structure according to the AF3 modelling.

Considering the variety in IgG responses towards different MSP2 variants, and the predicted structural heterogeneity, it’s possible that soluble, full MSP2 and soluble, truncated MSP2 are conformationally different and bind differently to the ELISA plate. We believe this indicates a significant change in conformation with removal of the conserved N and C terminus, which would mask the original targeted epitopes from IgG binding. This may also support the presence of structural heterogeneity within MSP2 variants even of the same family. A limitation of this study is the limited number of plasmas analyzed by peptide array. In the future, it would be interesting to expand our MSP2 peptide array platform to include more MSP2 variants, and more plasmas from varying exposure levels. This will elucidate the specific epitopes of MSP2 targeted by IgG from individuals with naturally acquired immunity. Importantly, we cannot show, experimentally, the structural heterogeneity of native MSP2 due to its disordered nature. However, protein structure predictions using AI-powered algorithms provide valuable insights on the antigenic nature of MSP2. The protein’s structure is a critical piece of information for continued research of MSP2 in the immunological context. Finally, a wider study of antigen sequence conservation is needed to address antigen polymorphism, which remains a challenge to the development of a strain-transcending malaria vaccine. Allele-specificity of the MSP2 immune response, and the rarity of IgG recognition of the conserved termini, supports the use of a multivalent vaccine including not only multiple different parasite proteins but also multiple variants of these proteins (68), which could emulate the effect of repeated infection. It is essential for our understanding of naturally acquired immunity, and for further vaccine design, to determine which epitopes of MSP2 are targeted by protective antibodies.

## Author contributions

JZ—data curation, formal analysis, investigation, visualization, writing—original draft and writing—review and editing; LM—data curation, formal analysis, investigation, visualization, and writing—original draft; MM, NSH, MR—investigation; PJ, NA—methodology, resources; BN, IR—resources; RS—investigation, resources; CS—resources; VY—resources, supervision; AF— funding acquisition, resources, writing—review and editing; DP— conceptualization, funding acquisition, project administration, supervision, writing—review and editing.

## Acknowledgments

We are grateful for the support provided by personnel involved in sample collection at the study site in Nyamisati and Karolinska University Hospital. The authors wish to acknowledge the support provided by the National Genomics Infrastructure (NGI)/Uppsala Genome Center and UPPMAX for their assistance with sequencing and computational infrastructure. The work conducted at NGI/Uppsala Genome Center was funded by RFI/VR and the Science for Life Laboratory, Sweden. Data processing and storage were made possible through resources supplied by the Swedish National Infrastructure for Computing (SNIC) at the Uppsala Multidisciplinary Center for Advanced Computational Science, with partial funding from the Swedish Research Council (VR 2018-05973). Additionally, bioinformatics analysis for long-read sequencing data and protein structure predictions in Alphafold2 were carried out using the Galaxy server, which is partially funded by the Collaborative Research Center 992 Medical Epigenetics (DFG grant SFB 992/1 2012) and the German Federal Ministry of Education and Research (BMBF grants 031 A538A/A538C RBC, 031L0101B/031L0101C de.NBI-epi, and 031L0106 de.STAIR [de.NBI])

## Data Availability Statement

The raw data supporting the conclusions of this article will be made available by the authors, without undue reservation.

