## Supplemental_information for "Naturally acquired IgG responses to *Plasmodium falciparum* do not target the conserved termini of the malaria vaccine candidate merozoite surface protein 2"

**Supplementary information**

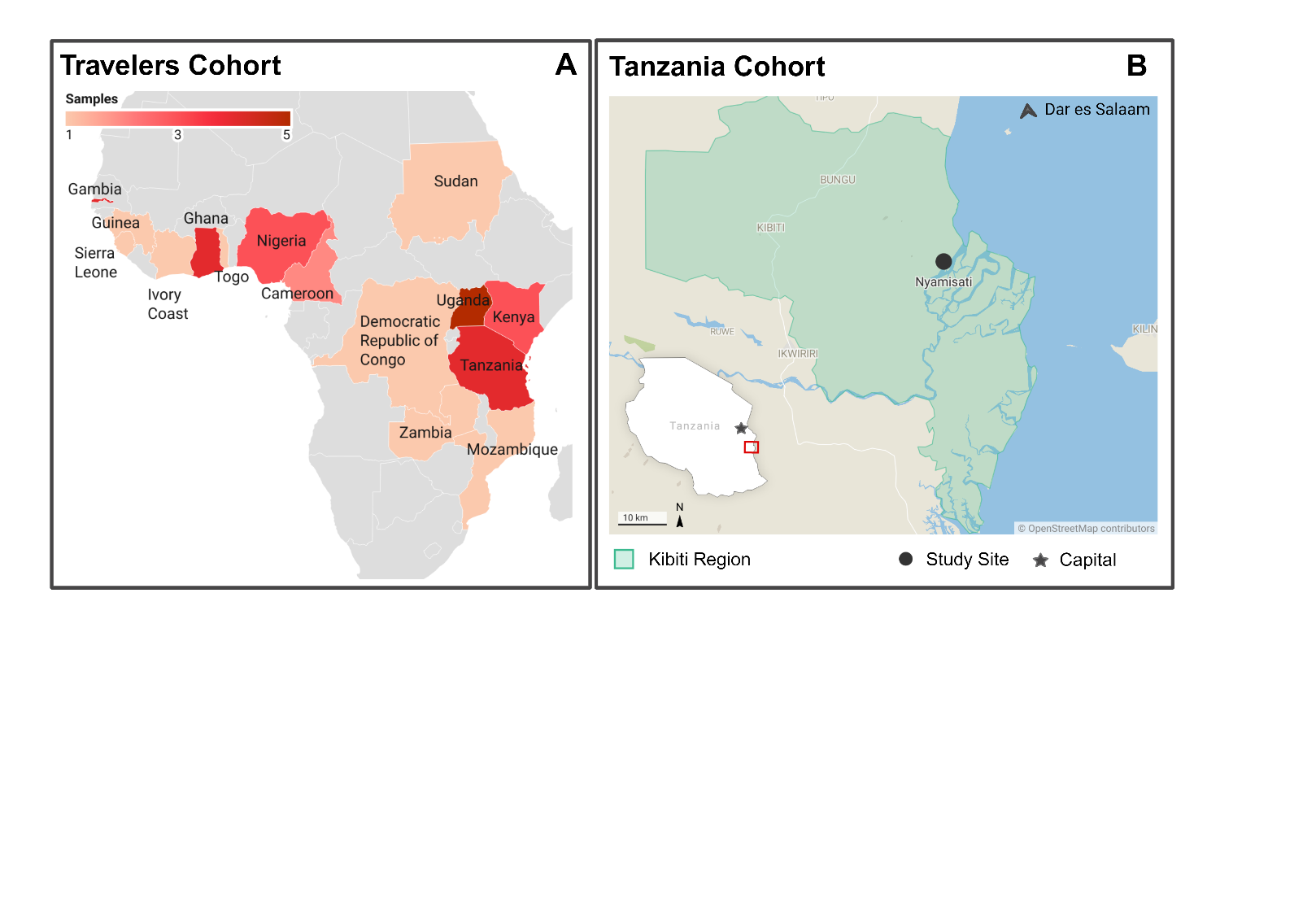
**Figure S1. Location of Nyamisati, Tanzania and reported country of infection for returning travelers with *Plasmodium falciparum* malaria.** (**A**) Travelers Cohort, where shades of red indicate the number of individuals returning from the country of infection (self-reported). (**B**) Location of Nyamisati within the United Republic of Tanzania. Figure created in Datawrapper.

**Table S1:** Primer and probe sequences used for *Plasmodium* species identification by qPCR, *msp2* amplification and barcoding for CCS, as well as size variant calling on *msp2* sequencing reads by *in silico* PCR.

| **Purpose and primer/probe name** | **Primer sequence (5’ → 3’)** |
| --- | --- |
| ***msp2* amplification and barcoding for CCS** **(Plaza *et al*, 2023)** | |
| 1^st^ reaction forward primer (msp2_fw) | ATGAAGGTAATTAAAACATTGTCTATTATA |
| 1st reaction reverse primer (msp2_rv) | TTATATGAATATGGCAAAAGATAAAACAA |
| 2^nd^ reaction forward primer | barcodes1001-to-1020-AATTTCTTTATTTTTGTTACC |
| 2^nd^ reaction reverse primer | barcodes1031-to-1050-GTGTTGCTGAAATTAAAAC |
| **Oligos for *msp2* family- and size variant-calling by *in silico* PCR** | |
| ICF1 | AGAAGTATGGCAGAAAGTAAGCCTCCTACT |
| ICF2 | AGAAGTATGGCAGAAAGTAATCCTCCTACT |
| ICF3 | AGAAGTATGGCAGAAAGTAAGCCTTCTACT |
| ICF4 | AGAAGTATGGCAGAAAGTAATCCTTCTACT |
| ICF_ML01 | AGAAGTATGGAAGAAAGTAATCCTCCTACT |
| ICF_GN01 | AGAAGTATGGCAGTAAGTAATCCTTCTACT |
| ICF_IT | AGAAGTATGACAGAAAGTAATCCTCCTACT |
| ICF_SD01 | AGAAGTATGTCAGAAAGTAAGCCTCCTACT |
| ICF_TG01 | AGAAGTATGACAGAAAGTAAGCCTCCTACT |
| ICF_GB4 | AGAAGTATGGCAGAAAGTAAGACTCCTACT |
| ICR | GATTGTAATTCGGGGGATTCAGTTTGTTCG |
| FC27F1 | AATACTAAGAGTGTAGGTGCAAATGCTCCA |
| FC27F2 | AATACTAAGAGTGTAGGTGCAGATGCTCCA |
| FC27F_KE01 | ACTACTAATAGTGTAGATGCAAATGCTCCA |
| FC27F_Dd2 | AATACTACTAGTGTAGGTGCAAATGCTCCA |
| FC27F_CD01_SN01 | AATACTAATAGTGTAGGTGCAGATGCTCCA |
| FC27F_HB3 | AATACTAAGAGTGTAGGTGCAAATGCTCCA |
| FC27R | TTTTATTTGGTGCATTGCCAGAACTTGAAC |

**Table S2:** **MSP2 variants sequenced in isolates from *P. falciparum*-positive travelers and expressed as recombinant proteins.** Protein sequences, translated from CCS reads, are shown in the native form (As expressed by the parasite). The sequences shown here include native signal peptides, GPI anchoring signals and predicted N-glycosylation sites.

| **Isolate (Reported Country of Origin)** | **Size variant (bp)** | **Reads / % Sample Coverage** | **Protein Sequence** |
| --- | --- | --- | --- |
| MSP2-FC27 1 (Gambia) | 292 | 25 / 72.0 | MKVIKTLSIINFFIFVTFNIKNESKYSNTFINN AYNMSIRRSMANEGSNTTSVGANAPNADT IANGSQSSTNSASTSTTNNGESQTTTPTAAD TPTATKSNSPSPPITTTKSNSPSPPITTTKSNS PSPPITTTESSSSGNAPNKTDGKGEESEKQN ELNESTEEGPKAPQEPQTAENENPAAPENK GTGQHGHMHGSRNNHPQNTSDSQKECTD GNKENCGAATSLLNNSSNIASINKFVVLISA TLVLSFAIFI |
| MSP2-IC 1 (South-East Asia) | 575 | 238 / 97.1 | MKVIKTLSIINFFIFVTFNIKNESKYSNTFINN AYNMSIRRSMEESNPSTGAGGSGSAGGSGS AGGSGSAGGSGSAGGSGSAGGSGSAGGSG SAGGSGSAGGSGSAGGSGSAGSGDGNGAN PGADAERSPSTPATTTTTTTTNDAEASTSTS SENPNHNNAETNPKGKGEVQKPNQANKET QNNSNVQQDSQTKSNVPPTQDADTKSPTA QPEQAENSAPTAEQTESPELQSAPENKGTG QHGHMHGSRNNHPQNTSDSQKECTDGNK ENCGAATSLLSNSSNIASINKFVVLISATLV LSFAIFI |
| MSP2-IC 2 (Ghana) | 707 | 77 / 80.5 | MKVIKTLSIINFFIFVTFNIKNESKYSNTFINN AYNMSIRRSMAESNPSTGAGGSGSAGGSA GGSAGGSAGGSAGGSAGGSAGGSAGGSAG GSAGGSAGGSAGGSAGGSAGGSAGGSAGG SAGGSAGGSAGGSAGGSAGGSAGGSAGGS AGGSAGGSAGGSAGSGDGNGANPGADAE GSSSTPATTTTTTTTTTTNDAEASTSTSSEN PKGKGEVQKPNQANKETQNNSNVQQDSQ TKSNVPRTQDADTKSPTAQPEQAENSAPTA EQTESPELQSAPENKGTGQHGHMHGSRNN HPQNTSDSQKECTDGNKENCGAATSLLNN SSNIASINKFVVLISATLVLSFAIFI |
| MSP2-FC27 2 (Kenya) | 376 | 571 / 95.1 | MKVIKTLSIINFFIFVTFNIKNESKYSNTFINN AYNMAIRRSMANKGSNTNSVGANAPNAD TIASGSQRSTNSASTSTTNNGESQTTTPTAA DTIASGSQRSTNSASTSTTNNGESQTTTPTA ADTIASGSQRSTNSASTSTTNNGESQTTTPT AADTPTATESSSSGNAPNKADGKGEESEKQ NELNESTEEGPKAPQEPQTAENENPAAPEN KGTGQHGHMHGSRNNHPQNTADSQKECT DGNKENCGAATSLLNNAANIASINKFVVLI SATLVLSFAIFI |

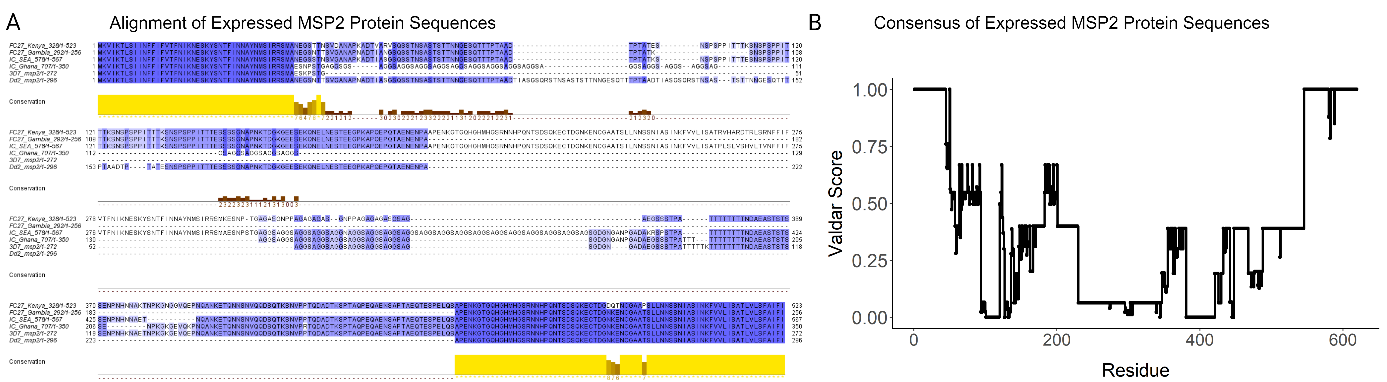

**Figure S2: Conservation and consensus of expressed recombinant MSP2 protein sequences.** MAFFT alignment of amino acid sequences for the native form of the four MSP2 variants engineered and recombinantly expressed for this study, in order: MSP2-FC27 2, MSP2-FC27 1, MSP2-IC 1, MSP2-IC 2 (**A**) in addition to two reference MSP2 sequences from 3D7 and Dd2 *P. falciparum* strains; colored by percentage identity. Normalized, Valdar consensus score of the four expressed protein sequences and two reference sequences (**B**). Alignment performed in Galaxy and visualized in Jalview.

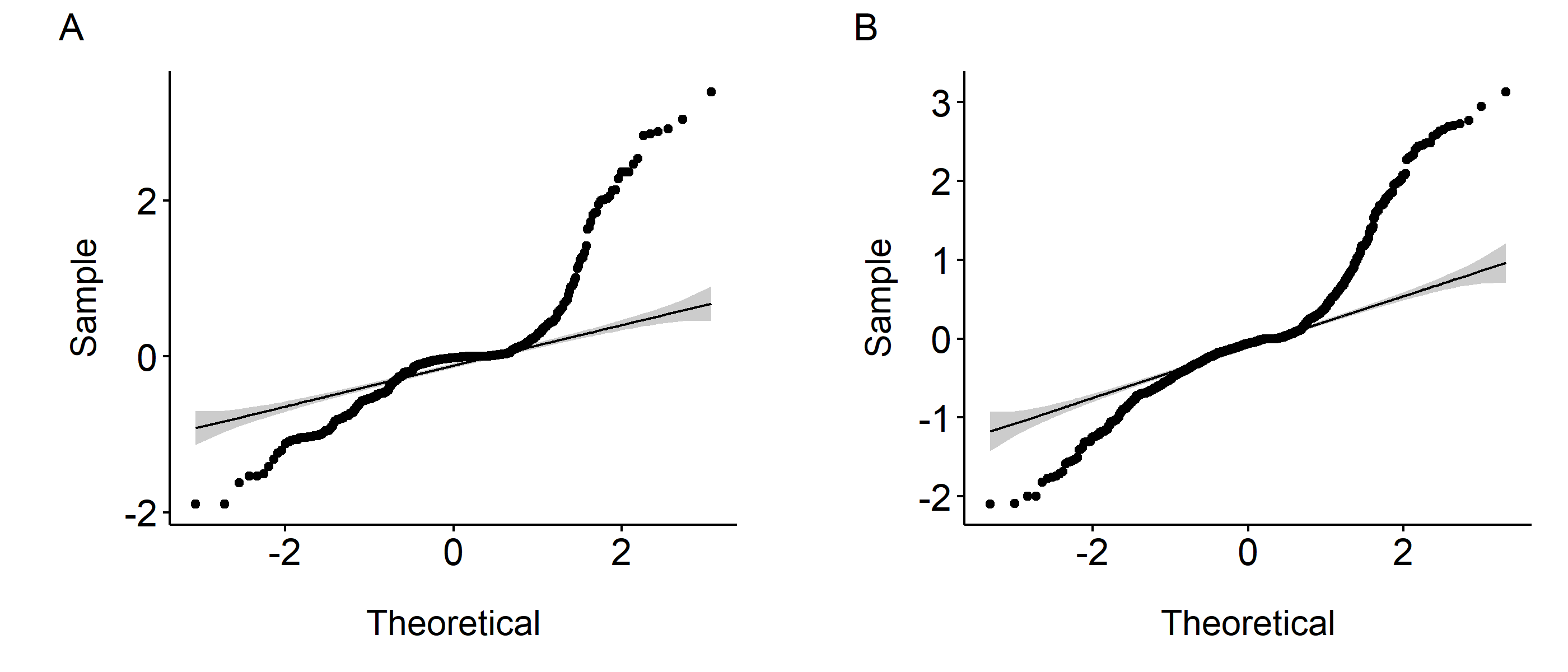

**Figure S3: Distribution of IgG responses by ELISA is non-normal**. QQ plot of residuals for IgG antibody responses in normalized OD, for Travelers (**A**) and Tanzania samples (**B**).

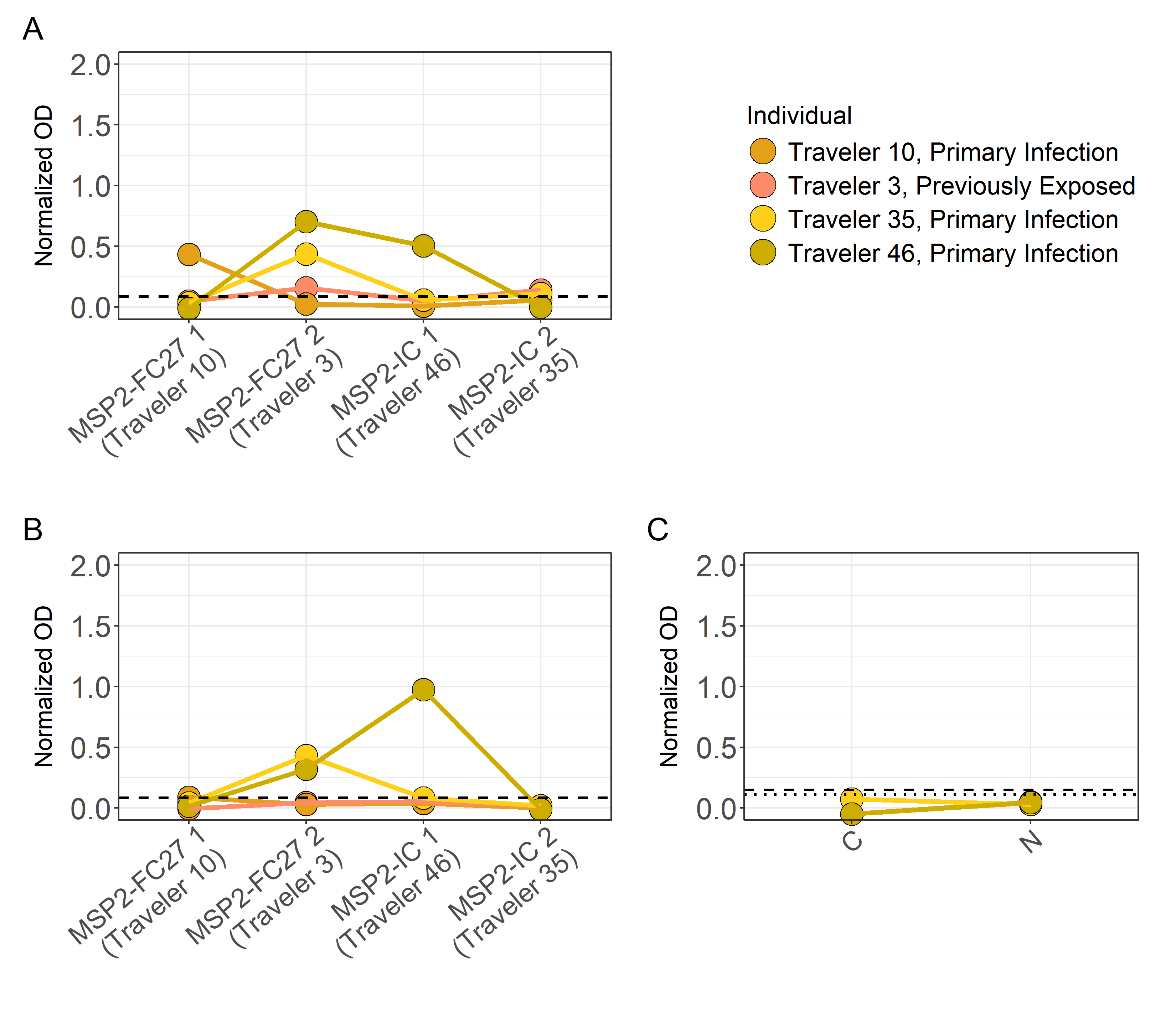

**Figure S4: IgG responses of autologous plasma towards MSP2 variants.** Plasma IgG from four travelers who were also carriers of the four expressed MSP2 variants used in this study, was evaluated for binding to the expressed full (**A**) or truncated (**B**) MSP2 variants, and conserved terminal peptides derived from the antigen (**C**). Dashed lines in (**A,B**) indicate mean cutoff for the shown variants. Dotted and dashed lines in (**C**) indicate mean cutoff for C- and N terminus peptides, respectively.

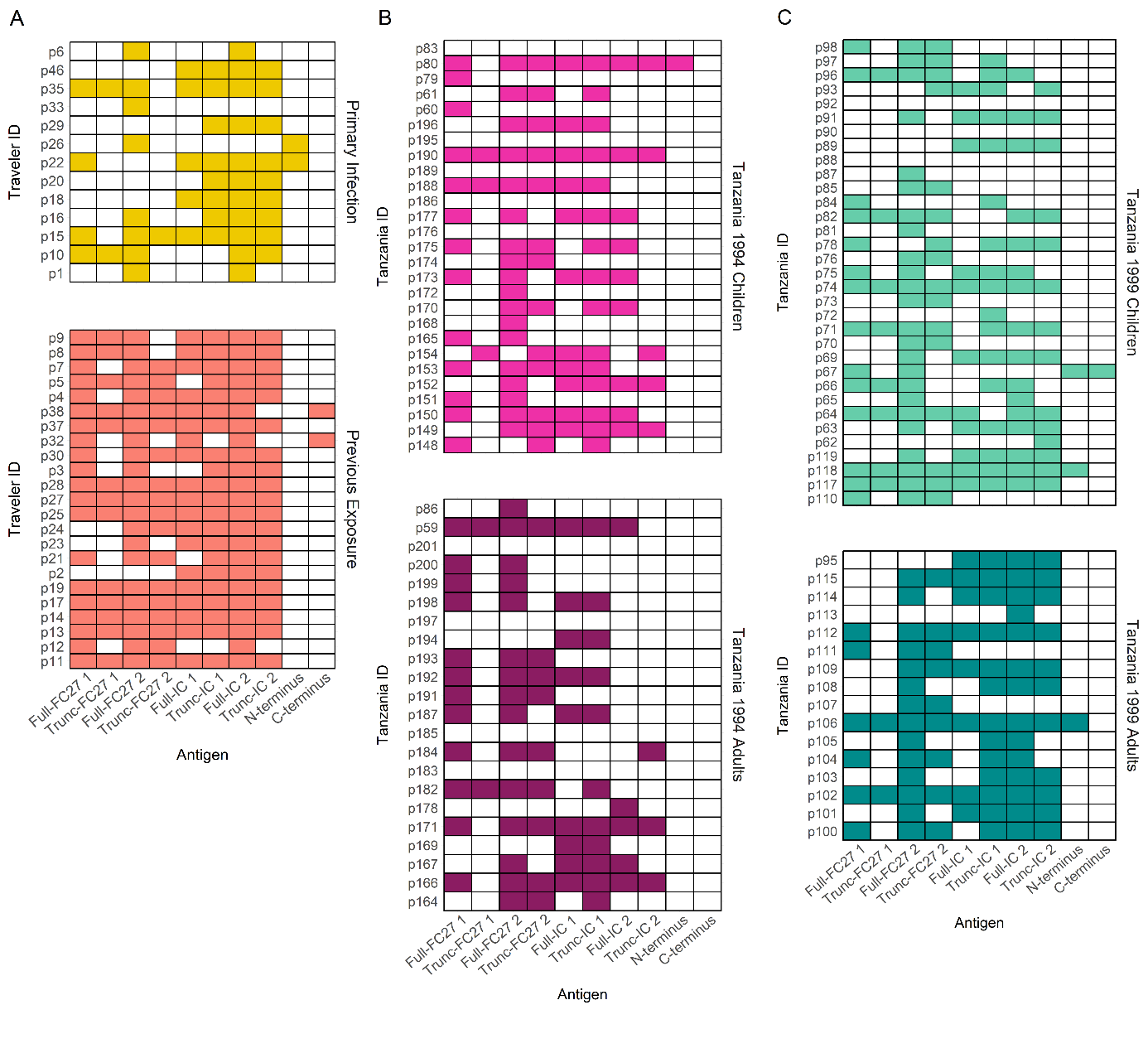

**Figure S5: Detected IgG binding of all tested plasmas to full and truncated versions of four MSP2 variants as well as peptides from the conserved N- and C- termini.** A binary response matrix was created after applying a negative threshold for each cohort and each of ten tested antigens, for primary infected and previously exposed Travelers (**A**), Tanzania 1994 children and adults (**B**), and Tanzania 1999 children and adults (**C**). Plasma samples whose IgG responses to a specific antigen passed the negative cutoff are indicated by colored tiles while those that did not pass the cutoff for the antigen are indicated by white tiles.

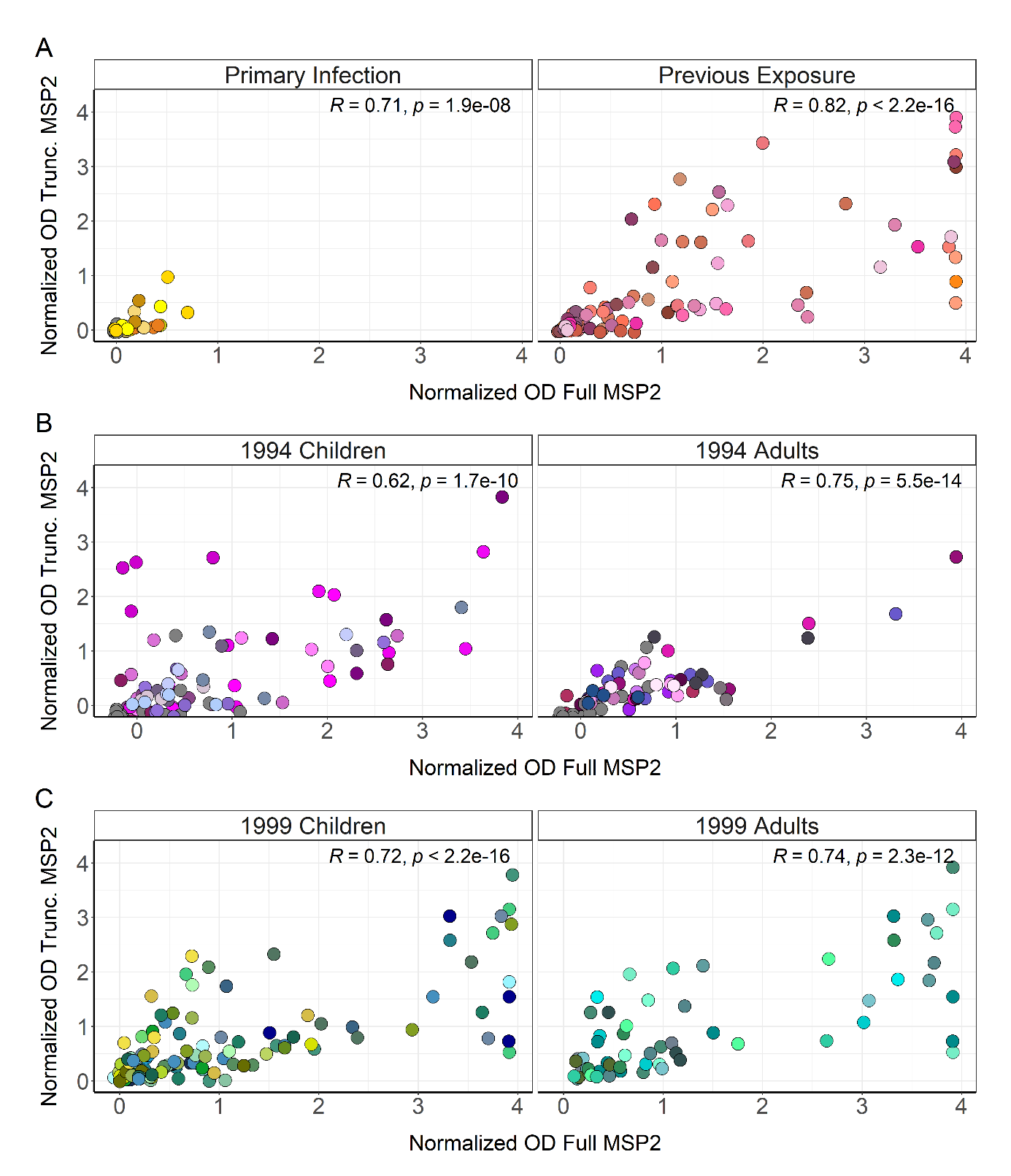

**Figure S6: Correlation of IgG responses to full and truncated MSP2 variants.** IgG responses in normalized OD values towards full MSP2 were plotted against those towards truncated MSP2 variants for primary infected and previously exposed travelers (**A**), Tanzania 1994 children and adults (**B**), and Tanzania 1999 children and adults (**C**). Correlation was estimated using Spearman rank correlation. Each color represents a different individual; therefore, every plot displays four data points of the same color corresponding to each of the four MSP2 variants tested.

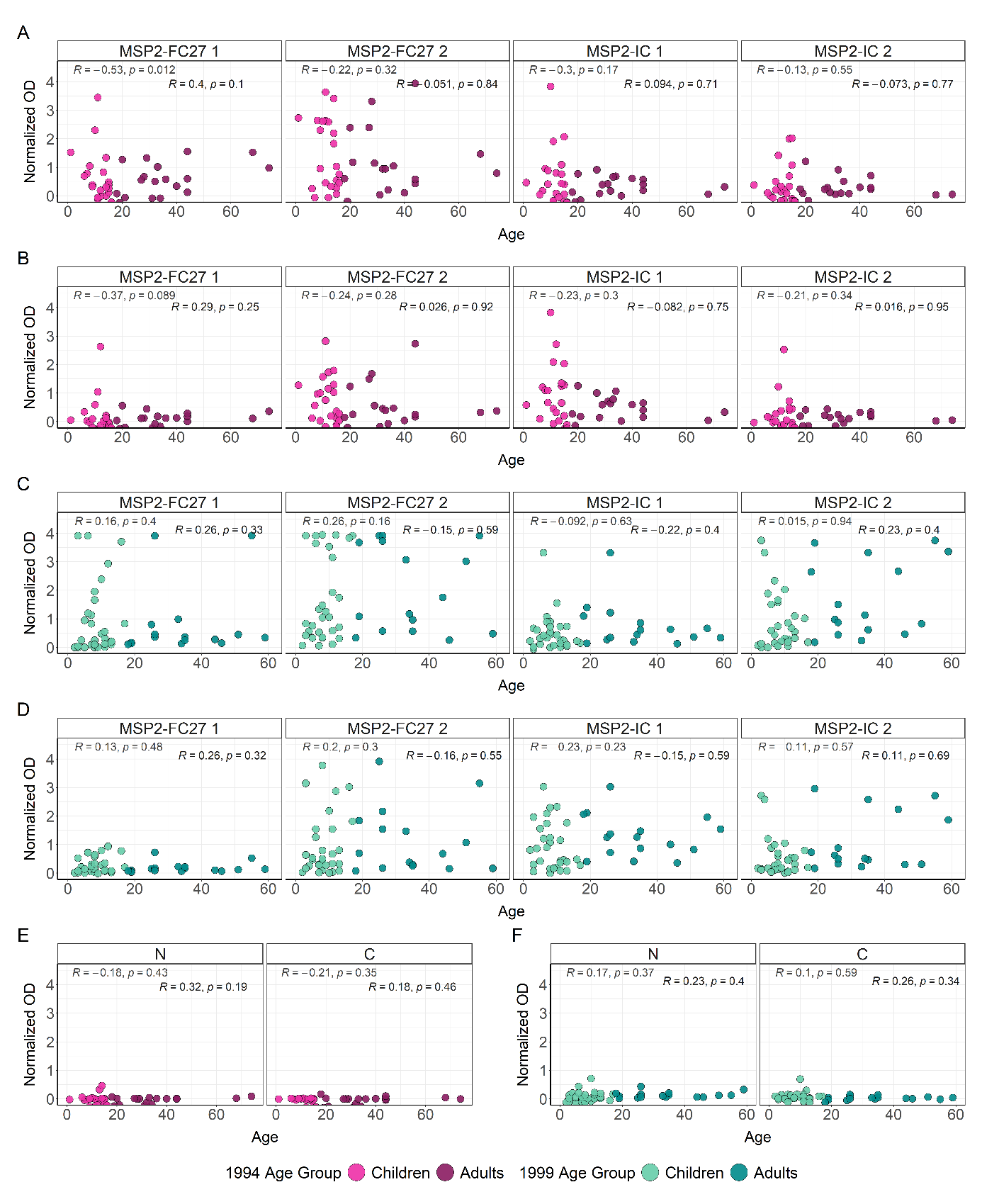

**Figure S7: IgG responses towards different MSP2 antigenic variants do not have a correlation with age.** IgG responses against full (**A**) and truncated (**B**) MSP2 variants and against conserved termini peptides (**E**), were plotted against age for individuals from Tanzania 1994. IgG responses against full (**C**) and truncated (**D**) MSP2 variants and against conserved termini peptides (**F**), were also plotted against age for individuals from Tanzania 1999. A Spearman rank correlation for each age group was fitted to sample data from all three sets. Sample data points are colored by survey and age group.

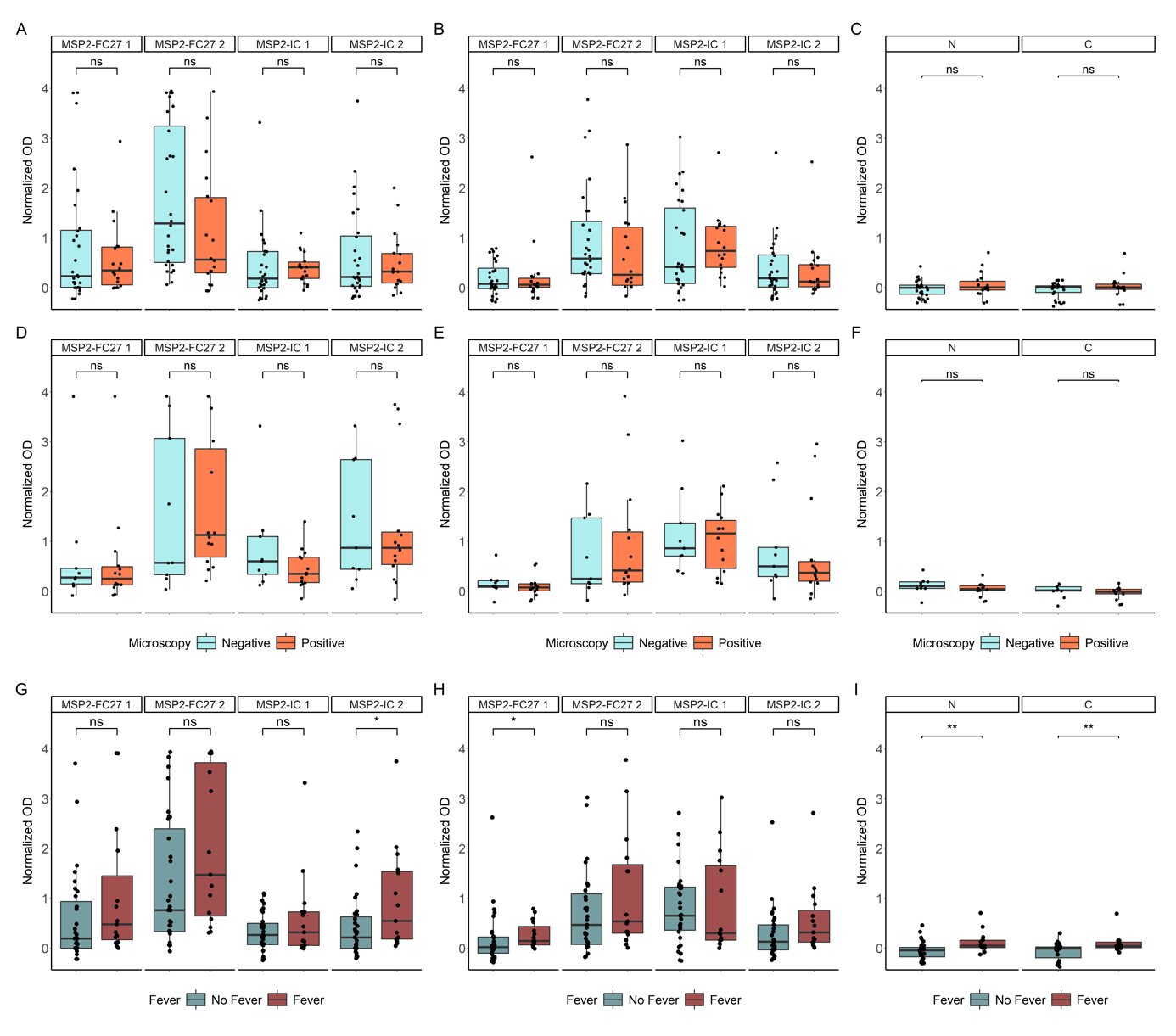

**Figure S8: IgG responses towards full and truncated recombinant constructs, as well as conserved MSP2 termini according to microscopy-positivity and fever status.** IgG responses towards full (**A**) or Truncated (**B**) constructs of MSP2, as well as towards conserved termini peptides (**C**) were compared for Tanzanian children who were microscopy-negative or microscopy-positive for *P. falciparum* at the time of sampling. IgG responses towards full (**D**) or truncated (**E**) constructs, and conserved termini peptides (**F**) were also compared for Tanzanian adults who were microscopy-negative or -positive. IgG responses towards full (**G**) or truncated (**H**) constructs, and conserved termini peptides (**I**) were also compared between Tanzanian individuals who presented in a febrile state or not at the time of sampling. No adults had recorded fever and were not included in the comparison. Statistical significance by unpaired Wilcoxon test (* < 0.05, ** < 0.01).

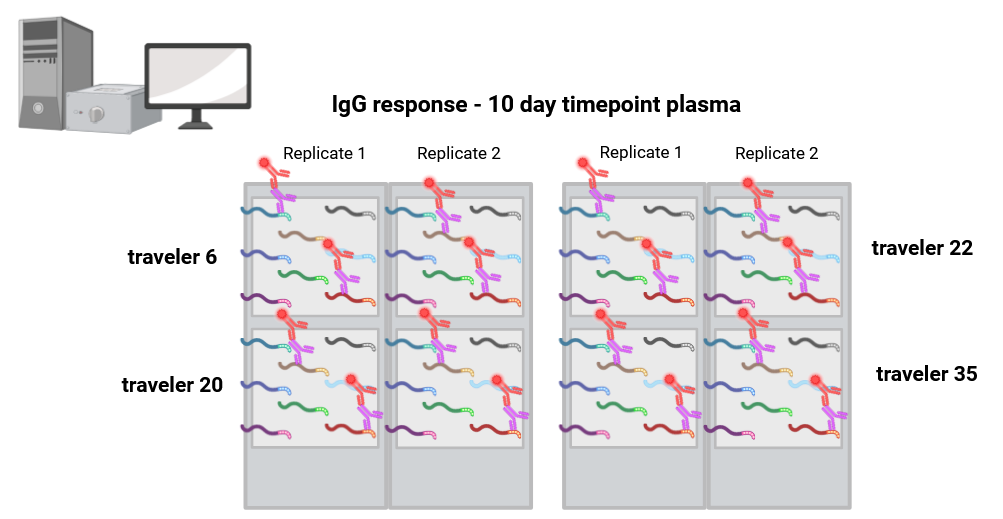
**Figure S9:** MSP2 peptide array setup schematic. The array consists of 128 13-mer MSP2 peptides, with 12 amino acid overlap giving single amino acid resolution. Included on each array were also positive control peptides from influenza and polioviruses. Figure created in BioRender.
